# Distinct ecotoxicological impairs induced by different *Microcystis* genotypes to Medaka fish

**DOI:** 10.1101/2024.12.24.630230

**Authors:** Benjamin Marie, Maiwenn Le Meur, Charlotte Duval, Manon Quiquand, Emilie Lance, Sébastien Duperron

## Abstract

One of the most prevalent and notorious bloom-forming freshwater cyanobacterial genus is *Microcystis*, whom toxicological impairs yet remain incompletely investigated. Based on our previous studies, we hypothesize that some emerging *Microcystis* metabolites, in addition to microcystins (MCs), are of (eco)toxicological concerns and should be further investigated. To this end, we explore the ecotoxicological potential of different *Microcystis* genotypes producing various bio-active metabolite cocktails, particularly cyanopeptides of different structural families including MCs, cyanopeptolins, microginins, anabaenopetins, aeruginosins or microcyclamides, on embryo/larvae and adult Medaka fish model.

Embryo and larvae exposures to the extracts of the four distinct *Microcystis* genotypes - comprising two MC-producing (PMC 728.11 and 807.12) and two non-MC-producing (PMC 810.12 and 826.12) strains – showed that PMC 810.12 and 826.12 respectively producing microginins, aeruginosins and microcyclamides exhibit early toxicity and teratogenicity, while PMC 728.11 presented rather larvae toxicity on hatched larvae, in agreement with its high MC content. In addition, we conducted a 4-days microcosm experiment with adult female Medaka exposed to environmental concentrations of these four *Microcystis* strain cultures to document the microbiome and metabolome responses. Fishes exposed to PMC 728.11 exhibited microbiota dysbiosis signature, while exposure to PMC 826.12 perturbated the fish digestion process inducing even more pronounced microbial and metabolic alterations. The two other strains provoked more moderate perturbations. These findings highlight toxic effects on fish exposed to both MC-as well as non-MC-producing cyanobacteria, suggesting complex interplay and effects of undocumented cyanobacterial bio-active and toxic compounds basides MCs during blooms of *Microcystis*.

## Introduction

One of the most common intense stress encountered by fauna in ponds and lakes is the occurrence of cyanobacterial blooms, which are now increasing in relation to anthropogenic effects (Huisman et al., 2018; Erratt et al., 2023). Despite their high productivity and quality as potential food sources (Cerri et al., 2021), cyanobacterial blooms in particular are not considered as a suitable source of animal feed because of associated adverse effects (Chen et al., 2021). Indeed, these proliferations may 1/ alter the aquatic trophic chain, 2/ lead to decrease in water dissolved oxygen (by consumption of their biomass by heterotroph bacteria), 2/ over-shadow other photosynthetic competitors, or 4/ directly threaten aquatic organisms through the production of deleterious compounds (Whitton et al., 2012). Bloom-forming freshwater cyanobacteria are well known producers of a wide range of bio-active metabolites, some of them being potentially toxic (Demay et al., 2019). To date, above 2,300 cyanobacterial compounds have been described from several genera, many of which are bioactive peptides, alkaloids, terpenes, macrolides and lipopolysaccharides (Jones et al., 2021). These products can be released into the water notably after bloom collapses, leading to several potential toxic effects to Human populations as well as on aquatic organisms (Codd et al., 2005).

Among the diversity of freshwater bloom-forming cyanobacteria, *Microcystis* represents one of the most proliferative genera worldwide, reported on all continents and on almost all countries so far (Harke et al., 2016). *Microcystis* dominates phytoplankton communities across a wide range of physico-chemical conditions. Several key functional traits may underpin the global success of *Microcystis* blooms. Thus, *Microcystis* blooms consist of diverse populations of distinct and co-proliferating clones/strains presenting various selective gene contents (Dick et al. 2021). Remarkably, several of the *Microcystis* genotypes can produce a multitude of bioactive accessory metabolites, including the hepatotoxins microcystins (MCs). Then, the persistence of their proliferation poses already-identified local risks to those using MC contaminated water resources for consumption, recreational activities, agriculture or fisheries (Sivonen and Jones, 1999; Codd et al., 2005). Moreover, up to 18 distinct accessory-metabolite biosynthetic gene clusters (BGCs) encoding non-ribosomal peptide synthase (NRPS) and/or polyketide synthase (PKS), together with other clusters encoding for ribosomally post-translationally modified peptide (RiPPs) or terpen producing enzymes have been detected within the various *Microcystis* published genomes (Cai et al., 2023). Several of these clusters encode enzymes involved in the biosynthesis of already known bioactive metabolites (such as microcystins, aeruginosins, cyanopeptolins, microginins, anabaenopeptins, cyanobactins and microviridins), whereas various of the remaining clusters seem to encode different enzymes responsible for the biosynthesis of yet-uncharacterized compounds (Humbert et al., 2013; Yancey et al., 2024). Moreover, different studies have already proposed that non-MC-producing *Microcystis* genotypes might also induce potential ecotoxicological impairs on aquatic organisms, in particular on fish, key contributors of aquatic ecosystem functioning (Marie, 2020; de Almeida et al., 2024; Banerjee et al., 2021).

One of the most well-known cyanotoxins frequently produced by *Microcystis* is MC-LR, a hepatotoxin that mainly accumulates in the liver with deleterious consequences for fish physiology and reproductive processes even at low chronic doses (Qiao et al., 2016; Qiao et al., 2019). However, Saraf and coworkers (2018) have shown on zebrafish embryos that the MCs alone did not explain all toxicological effects induced by various *Microcystis* strains, and that other molecules may provoke cumulative or synergic deleterious effects. Also, yellow-tail tetra embryos showed developmental impairs when exposed to *Microcystis* biomasses regardless of whether the cultured strains were producing MCs or not (Fernandes et al., 2019). A step further, similar works performed with non-MC-producing *Microcystis* strains also showed high embryonic toxicity of extracts containing microginins and cyanopeptolins (de Almeida Torres et al., 2023). Purified microcylamide extracted from a *Microcystis* strain has also been tested on zebrafish embryos showing a toxic concentration range similar to that of pure MC-LR (de Freitas et al., 2024). Interestingly, several studies attempt to compare the tissular, physiological and molecular effects induced by both MC-producing and non-MC-producing *Microcystis* strains, showing that the latter ones provoked intense but distinct perturbation on the organs of adult fish models (Le Manach et al., 2018; Sotton et al., 2017). Recently, non-microcystin-producing *Microcystis* strains have also been shown to impair cell line viability and reproductive gene expression (Harshaw et al., 2024).

Considering that freshwater fish are overall exposed to cyanobacteria and their toxins through oral ingestion, the response of gut-associated microbiota has also been investigated as a potential primary target for microbiome-aware ecotoxicological concerns linked with *Microcystis* proliferations (Duperron et al., 2020; Duan et al., 2020; Gallet al., 2023). Recent works also focussed on fish appetite depletion in the presence of *Microcystis* biomass, showing subsequent neuro-endocrine alteration (Nui et al., 2024; Long et al., 2024). In parallel, other smaller molecules, such as terpenes, could also present threaten to the fish as teratogenicity (Jonas et al., 2015). Such effects might be related to retinoid production highlighted in biomass extracts of lab-cultured cyanobacteria *M. aeruginosa* (Jonas et al., 2015). Altogether, we thus hypothesize that the chemical variety of bioactive metabolites produced by the diverse *Microcystis* genotypes may be of greater ecotoxicological concern than the, so-called cyanotoxins, MCs and needs to be further investigated (Janssen, 2019).

The present study seeks to gain insight into the diversity of the toxic mechanisms of selected *Microcystis* strains belonging to distinct genotypes regarding toxicological effects on the embryos and the adults of the Medaka fish *Oryzias latipes*. Four selected strains were previously characterized to identify the potentially bioactive metabolites they produce using combined genome mining and metabolomic investigations. Based on the implementation of a comprehensive battery of bioassays with responses measured at the individual (*e.g.*, mortality, developmental abnormalities) and sub-individual (e.g., molecular, microbiota effects) levels, this study provides new insights into the complexity of toxic potencies of four different *Microcystis* genotypes producing, or not, MC and other cyanopeptide families. Overall, this work reveals the diversity of effects and large spectra of alterations they may induce on fish fitness.

## Material and methods

### *Microcystis* strain materials

The four *Microcystis* strains PMC 728.11, PMC 807.12, PMC 810.12 and PMC 826.12 were obtained from the Paris Muséum Collection. The PMC 728.11 strain was collected and isolated from Lake Juanon around Valence (France) (Halary et al., 2020), whereas the PMC 807.12, PMC 810.12 and PMC 826.12 strains were isolated from a recreational pond in Champs-Sur-Marne (Ile-de-France, France) (Gaetan et al., 2023). These four French strains were specifically selected according to the proximity of their geographic origin, because they belong to diverse clades of the global *Microcystis* phylogeny, and produce different sets of bio-active metabolites, as further described.

### *Microcystis* genome analysis

We first downloaded all available *Microcystis* genomes from the NCBI database and then removed low-quality or duplicated ones. Publicly available genome assemblies belonging to the *Microcystis* genus were retrieved from the NCBI FTP server using both genus name identifiers of 1125 [*Microcystis*] on 14 January 2022. In total 344 non-redundant assemblies were collected from GenBank and/or RefSeq databases. Genome completeness was then assessed using CheckM (Parks et al., 2015) to selected high quality genomes (completeness >95% or contamination <5%). From this 278-*Microcystis* genome dataset, a core genome consisting of 80 single-copy genes shared by all genomes was retrieved using Roary (sequence similarity threshold of 90%, multiple alignment performed by MAFFT) (Page et al., 2015; Katoh and Standley, 2013) based on coding gene prediction carried out by Prokka (Seemann 2014). The obtained concatenated core gene sequences multiple alignment of 686,669 nucleotides was used to infer a phylogenetic tree using RaxML v8.2.12 (Stamatakis et al., 2012) (GTR-Gamma model, 100 bootstraps). The phylogenetic tree was visualized and annotated using iTOL (Letunic and Bork, 2021), with *M. aeruginosa* Ma_SC_T_19800800_S464 chosen as the outgroup according to Cai et al. (2023).

Putative byosinthetic gene clusters (BGCs) encoded within the four *Microcystis* genomes presently investigated were then run through AntiSMASH 6.0 (Blin et al., 2021) with detection strictness set to “relaxed” and all extra features activated. BGC matches were further explored if they contained NRPS (cyanopeptolin, aeruginosin and anabaenopetin), NRPS/PKS (microcystin and microginin), PKS (MAA) or RiPP (aerucylamide or microviridin).

### *Microcystis* biomass production, metabolite extraction and toxicological tests on Medaka fish embryos and larvae

The four *Microcystis* strains were cultured non-axenic in BG-11 medium (Stanier et al. 1979) at 22°C in 250-mL Erlenmayer’s vessel, with a photon flux density of 12 µmol.m^-2^.s^-1^ and a 13:11h light:dark cycle. Cellular biomasses were collected by centrifugation (3,200 g, 30 min at 15°C) and intracellular metabolites were extracted from one gram of freeze-dried biomass with 10 mL of methanol:water (70:30) with 0.1% of formic acid solution. Cell lysis was performed by sonication: 3 cycles of 30 s, 10 s break between each cycle, at 80% of the maximum intensity (SONICS Vibra Cell, Newton, CT, USA; 130 Watts, 20 kHz). Samples were then centrifuged (10 min, 13,400 g, 4°C), and the supernatants were collected and stored in the dark at -20°C prior to the exposure of embryo and larvae of Medaka fish and additional mass spectrometry characterization. Whole MC concentrations in the extracts (sum of all variants) were estimated according to LC-MS/MS analysis (Olokotum et al., 2022).

All fish studies were carried out on *Oryzias latipes* (Cab lineage) maintained in the Medaka fish facility of the Muséum National d’Histoire Naturelle (MNHN - Paris, France), under standard controlled conditions and in accordance with relevant European animal protection law. Above two hundreds of 6-month-old adults were maintained in reproductive regime (25±1°C and under a photoperiod of 15h/9h light/dark) in 6-L tanks by groups of ten individuals (5 females and 5 males) in order to induce reproduction and to daily collect manually fertilized eggs on female abdomen. Tanks were connected to a recirculating flow-through system supplied with a mixture of tap and deionized water (1:2). Fish were fed three times a day with granulate of GEMMA MICRO ZF-300µm (Proteins 59%; Lipids 14%; Fiber 0,2%; Minerals 14%; Phosphorus 1,3%; Calcium 1,5%; Sodium 0,7%; Vitamin A 23000 UI. Kg^-1^; Vitamin D3 2800 UI.kg^-1^; Vitamin C 1000 mg.kg^-1^; Vitamin E 400 mg.kg^-1^; Skretting, Norway). The eggs were collected in the evening, approximately 1-2 h post-fertilization, were rinsed and kept in Yamamoto’s 1X medium (0.128 M NaCl; 0.27 mM KCl; 0.14 mM CaCl_2_; 0.24 mM NaHCO_3_) for the rest of the experimentations performed at 25±1°C and under a photoperiod of 15h/9h light/dark (Yamamoto, 1956).

First, a part of the freshly collected eggs were incubated in 24-well exposure plates (4 embryos per well in 1 mL, 5 replicates, for a total of 20 embryos per condition) with extracts of the different *Microcystis* strains diluted at the same maximal concentration of 14 mg of extracted *Microcystis* cells per mL in 1% methanol (final concentration). A control condition (methanol solvent used for metabolite extraction) was performed as negative control on each experimental plate. Medium containing the different extracts was completely replaced after 24h incubation, then the survival, the development arrest and the occurrence of head or tail malformation were individually observed after 48h exposure (Supplementary figure S1). Notice that acceptably low larvae mortality (<10%) occurred with the control condition in accordance with OCDE 236 test guidelines (OCDE 2013).

Also, a fraction of the freshly fertilized embryos was maintained in 6-cm Petri dishes filled with Yamamoto’s medium renewed daily until the egg hatching (8 dpf), then larvae were collected and incubated 24h in 12-well exposure plates (3 embryos per well in 3 mL, 4 replicates, for a total of 12 embryos per condition) with extracts of the different *Microcystis* strains diluted at the same maximal concentration of 14 mg of extracted *Microcystis* cells per mL in 1% methanol (final concentration). A control condition (methanol solvent used for metabolite extraction) was performed as negative control on each experimental plate. Embryos were then monitored and the larvae survival was measured after 24h of incubation. Notice that no larvae mortality occurred with the control condition (Supplementary figure S1).

At the end of the procedure, all exposed embryos and larvae were euthanized by overdose of MS-222 (CAS n°886-86-2, Sigma Aldrich). The whole procedure was performed twice and the data combined, as presenting no significant variation.

### Exposure of adult female Medaka to four different *Microcystis* cultures

Experimental procedures were carried out in accordance with European legislation on animal experimentation (European Union Directive 2010/63/EU) and were approved for ethical contentment by an independent ethical council (CEEA Cuvier n°68) and authorized by the French government under reference number APAFiS#19316-2019032913284201 v1.

Balneation fish experiments were performed in 10-L aquaria, each containing 4 specimens of 6-months old adult female medaka (Supplementary figure S2). Aquaria water was stabilized under constant aeration (bubbling devices) for 1 month, and fishes acclimatized for 1 week prior to exposure. Water parameters were monitored daily (pH, temperature, conductivity, nitrates and nitrites), faeces were removed daily by aspiration, and half of the water was replaced with freshwater (2/3 osmosis (RiOs 5, Merck Millipore) and 1/3 filtered). Fish were exposed to constant temperature (23.1 ±0.7 °C), pH (6.7 ±0.2) and conductivity (223 ±6.8 *µ*S.cm^-1^), to low levels of nitrates and nitrites (≤ 0.05 mg.L^-1^) to a 12h:12h light/dark cycle. They were fed twice daily (∼3-5% of the fish biomass per day) with GEMMA MICRO ZF-300µm.

Fish were then exposed for 4 days to water or water containing an environmentally relevant concentration of a culture of *Microcystis* PMC 728.11, PMC 807.12, PMC 810.12 and PMC 826.12 strains (100 *µ*g.L^-1^ Chl *a*) (Foucault et al., 2022). All *Microcystis* strain cultures were initially grown in Z8 medium at 25 ±1 °C with a 16h:8h light/dark cycle (at 14 *µ*mol.m^-2^.s^-1^) in 2-L bottles. The initial concentrations were estimated by performing Chlorophyll *a* extractions and absorbance measurements as a proxy using a spectrophotometer (Cary 60 UV-Vis, Agilent). At the beginning of the exposure, adapted volumes of each *Microcystis* culturewere added to the aquarium to reach the initial concentration of 100 *µ*g.L^-1^ Chl *a*, anticipated to induce noticeable effects according to previous experiments performed by Gallet et al. (2023), and the same volume was added at day two (after 48h) to maintain exposure level. Then, four fish per aquarium were sampled on day 4 (96h), as well as water tank and faeces samples, that were pooled in individual sample for each aquarium.

Fish were anesthetized in 0.1% tricaine methane sulfonate (MS-222; Sigma, St. Louis, MO) buffered with 0.1% NaHCO_3_ and sacrificed. Whole guts, muscles and livers were dissected, flash-frozen in liquid nitrogen and stored at -80 °C. Aquarium water samples were filtered on a 0.22-*µ*m filter (Nucleopore Track-Etch Membrane) and frozen. *Microcystis* cultures (10 mL) were centrifugated (10 min, 10 °C, 3,220 g) and pellets were frozen. Faeces pellets were directly frozen and stored at -80 °C.

### Fish metabolite extraction

All samples were suspended in cold extraction solvent, consisting of 75-25% methanol-water, 1 mL per 100 mg of fresh tissue on ice (Colas et al., 2020). A prior mechanical homogenization (GLH850 OMNI; 25 000 r. min^-1^; 30s) was performed on all sample types including fish guts, livers, muscles and faeces, except tank water. Samples were then sonicated (Sonics Vibra-Cell VCX 13; 60% amplitude; 30s, x3) and centrifugated (10 min; 4°C; 15,300 g), the supernatant containing metabolite extracts were collected and kept at -20°C prior to subsequent metabolomic investigation by mass spectrometry. The remaining extraction pellets were kept at -80 °C and further used for subsequent DNA extraction of guts, faeces and tank waters (Duperron et al., 2023).

### Mass spectrometry data processing and analysis

The metabolite composition from *Microcystis* extracts and cultures, fish livers, guts muscles and faeces and tank water were analysed by ultra-high-performance liquid chromatography (UHPLC; ELUTE, Bruker) coupled with a high-resolution mass spectrometer (ESI-Qq-TOF Compact, Bruker). For each sample, 2 μL were injected, and molecule separation was performed by a Polar Advance II 2.5 pore C_18_ (Thermo Fisher Scientific, Waltham, MA, USA) chromatographic column at a flow rate of 300 μL.min^−1^ under a linear gradient of acetonitrile acidified with 0.1% formic acid (from 5 to 90% in 15 min). The electrospray ionization (ESI) system was calibrated with capillary temperature was set to 200°C, the source voltage was 3.5 kV, and the gas flowrates were 8 L.min^-1^. Then, ions were analyzed in the range 50–1500 *m/z* performed by collision ion dissociation (CID) and autoMS/MS in positive or negative modes with information dependent acquisition (IDA). Thus, compounds were analyzed in simple MS positive modes without quadrupole fragmentation at 2 Hz frequency acquisition rate, then top-intensity ions (> 5,000 counts in single MS (MS1)) were individually selected by the quadrupole in a window of 10 Da and fragmented in a collision cell (MS2) with a selective exclusion window of 30s (except for ions presenting a count-intensity increase superior to 3 times) with selective ion collision set between according to ion intensity and mass (10-50 eV ramp; 50/50% time window; 2-8 Hz adapted according to ion count intensity), in consecutive cycle times of 2.5s. The resulting ions of the fragmentation of their respective parent were transferred and detected.

MS data were processed using MetaboScape 4.0 software (Bruker, Bremen, Germany) for recalibration of each sample analysis (<1ppm, according to acetate formate internal standard), peak detection and selection of ions whose intensity was greater than 5,000 counts in at least 10% of the set of samples and minimal RT-correlation coefficient set at 0.7. Furthermore, different states of charge (1+, 2+ and 3+ or 1-, 2- and 3-) and classical adducts were grouped together and the “area-under-the-peak” was determined in order to generate a unique global data matrix containing semi-quantification results for each metabolite in all analyzed samples’ peak for each analyte (characterized by the respective mean mass of its neutral form and its corresponding retention time). Data QC and Blank samples (injected every 6 samples and every triplicats, respectively) were examined in order to ensure the reproducibility and robustness of the whole data series. First analyte annotations were attempted according to respective ion mass and isotopic pattern by automatic match with the CyanometDB 2.0 database which contains above 2,300 chemical formulas of already known specialized metabolite produced by cyanobacteria (Jones et al., 2021) and the natural product NPAtlas 2.0 database (van Santen et al., 2022). The files containing all the MS2 fragmentation information for each ion analyzed were exported in mgf format using MetaboScape 4.0 (Bruker, Bremen, Germany) software before being used for the generation of the molecular network of structural similarity using GNPS algorithm (Wang et al., 2016) using the MetGem software (version 1.3.6) (Olivon et al., 2018). Additional metabolite annotations were also made from MS2 data by comparing of fragmentation profiles with GNPS algorithm (with parameters set at: *m/z* tolerance=0.02 Da; minimum matched peaks=4; topK=10; minimal cosine score=0.70) using GNPS, NIH, MS-DIAL and EMBL public and generalist spectral databases.

### DNA extraction, bacterial 16S rRNA gene sequencing and analyses

DNA was extracted from the pellets obtained after metabolites extraction using the DNeasy® PowerLyser® PowerSoil® Kit (Cat N° 12855-100, Qiagen, Germany following manufacturer’s protocol. Mechanical lysis was carried out with a FastPrep-24^TM^ 5G BeadBeater (MP Biomedical, USA) for 5 cycles of 30 s ON, 30 s OFF at a frequency of six beats per second. (Gallet et al., 2023). An extraction-blank control sample was also performed.

The V3-V4 region of the 16S rRNA-encoding gene was amplified using primers 341F (5’-CCTACGGGNGGCWGCAG -3’) and 806R (5’-GGACTACVSGGGTATCTAAT-3’) using the following program: initial denaturation (94°C, 3 min); 35 cycles (94°C, 45 s; 55°C, 60 s; 72°C, 90 s); elongation step (72°C, 10 min). Products were sequenced on an Illumina MiSeq 250×2 bp platform (GenoToul, France). Reads were deposited into the Sequence Read Archive (SRA) database (accession number PRJNAxxx (samples SRRxxx - SRRxxxx). Paired-end reads were demultiplexed, quality controlled, trimmed and assembled with FLASH (Hahaut and Picelli 2022). Sequence analysis was performed using the QIIME 2 2020.11 pipeline (Bolyen et al. 2018). Chimeras were removed and sequences were trimmed to 360 pb then denoised using the *DADA2* plugin, resulting in Amplicon Sequence Variants (ASVs). ASVs were affiliated based on the SILVA database release 138 (Quast et al., 2013) using the *feature-classifier* plugin and *classify-sklearn* module (Pedregosa et al., 2011; Bokulich et al., 2018). Sequences assigned to Eukaryota, Archaea, Mitochondria, Chloroplast and Unassigned were removed from the dataset, then libraries from each sample were rarefied to 6,282 reads.

### Data treatment and statistical analyses

For the metabolome dataset, the MetaboAnalyst 5.0 platform (Pang et al., 2021) was used to perform data matrix normalization of metabolomic data according to *Pareto* and mean-centered and divided by the standard deviation for the different metabolite matrices. The strain metabolite correlation was performed according to Spearman rank calculation and visualized on a heatmap with hierarchical classification using MetaboAnalyst 5.0. A clade based on the presence or absence of coding gene sequences was obtained with a dendrogram (according to Jaccard’s similarity/dissimilarity index, Ward reconstruction, and Silhouette analysis to find the optimal number of clusters) using R software (version 1.4.1103).

For the microbiome dataset, alpha- and beta-diversity analyses were performed using the MicrobiomeAnalysis 2.0 online platform (Lu et al., 2023), Principal coordinates analyses (PCoA) based on Bray-Curtis distances were performed to examine the dissimilarity of bacterial community compositions among groups. Among- and within-group variance levels were compared using PERMANOVA (999 permutations) and Pairwise Adonis (999 permutations), respectively. Differentially-abundant taxa across groups were identified using the linear discriminant analysis (LDA) effect size (LEfSe) tool (Chang et al., 2022).

Considering the statistic robustness according to the fact that all microbiome and metabolomic investigations have been developed on the different organs of the same individuals, multi-omics integration of the data has been performed (Marie, 2020). To this end, multi-block sPLS-DA performed using DIABLO tool (Singh et al., 2019) offers the opportunity to depict the whole molecular network of correlation within the explored dataset. The integration of metabolome and microbiome datasets (area-under-the- and ASV counts in the same samples) was performed using the *mixOmics* package in R (Rohart et al., 2017). Pareto scaling was applied on the metabolome data and a centred log-ratio transformation then a pre-filtering keeping only abundant ASVs, (*i.e.* representing at least 1% of the reads in at least one sample), was applied on the microbiome data. A correlative analysis was performed with DIABLO (Singh et al., 2019), a multi-omics framework created by the MixOmics team to analyze putative correlated dynamics between gut microbiota ASVs and gut metabolites. Briefly, the *block.plsda()* function integrates several datasets (named “blocks”) and performs a Pattern Latent Structure Discriminant Analysis. A correlation score for the given blocks was computed with the *plotDiablo()* function. Then, relevance networks displaying the most discriminant covariates (metabolites, ASVs) were produced using the *network* function with Pearson correlation cut-offs.

## Results and discussion

### Diversity of the four selected *Microcystis* strains

Among the monoclonal *Microcystis* strains available in live strains collections, we selected four strains representative of the diversity of *Microcystis*, considering genotypes that can be co-proliferating in the same bloom (Figure 1). Under photonic microscope (Figure 1A), both monoclonal strains cultivated under non-axenic conditions in Z8 media (Hamlaoui et al, 2022) exhibit individualized cell development that have been identified as both corresponding to *Microcystis aeruginosa* species, without showing discriminative characters. However, the different morpho-species of *Microcystis* spp. have recently been revealed to belong to a single cosmopolitan genus that comprises various genotypes discriminated at the genome level, without correspondence to any of the previously described diversity in colony shape or morpho-species (Cai et al., 2023). A genomic investigation of the respective phylogenetic position of these four strains clearly shows they belong to four distinct clades, evenly spread over the *Microcystis* phylogenomic tree (Figure 1B). A detailed investigation of the cortege of accessory and potentially bio-active, if not toxic, metabolites produced by these respective strains reveals a large chemical diversity (supplementary table S1-4). Indeed, both genome mining and direct metabolomic investigations reveal the production of diverse metabolite families, with for example PMC 728.11 and PMC 807.12 both producing MCs, whereas both PMC 810.12 and PMC 826.12 produce microginins and microcyclamides, among various other metabolite families (Fig. 1C; supplementary figure S3). Taken together, these data exquisitely illustrate how chemically diverse *Microcystis* genotypes are (Humbert et al., 2013; Le Manach et al., 2019). This result urges us to fully consider the occurrence of a broad intra-species diversity when investigating the (eco-)toxicological risk associated with *Microcystis* blooms.

**Figure 1.**
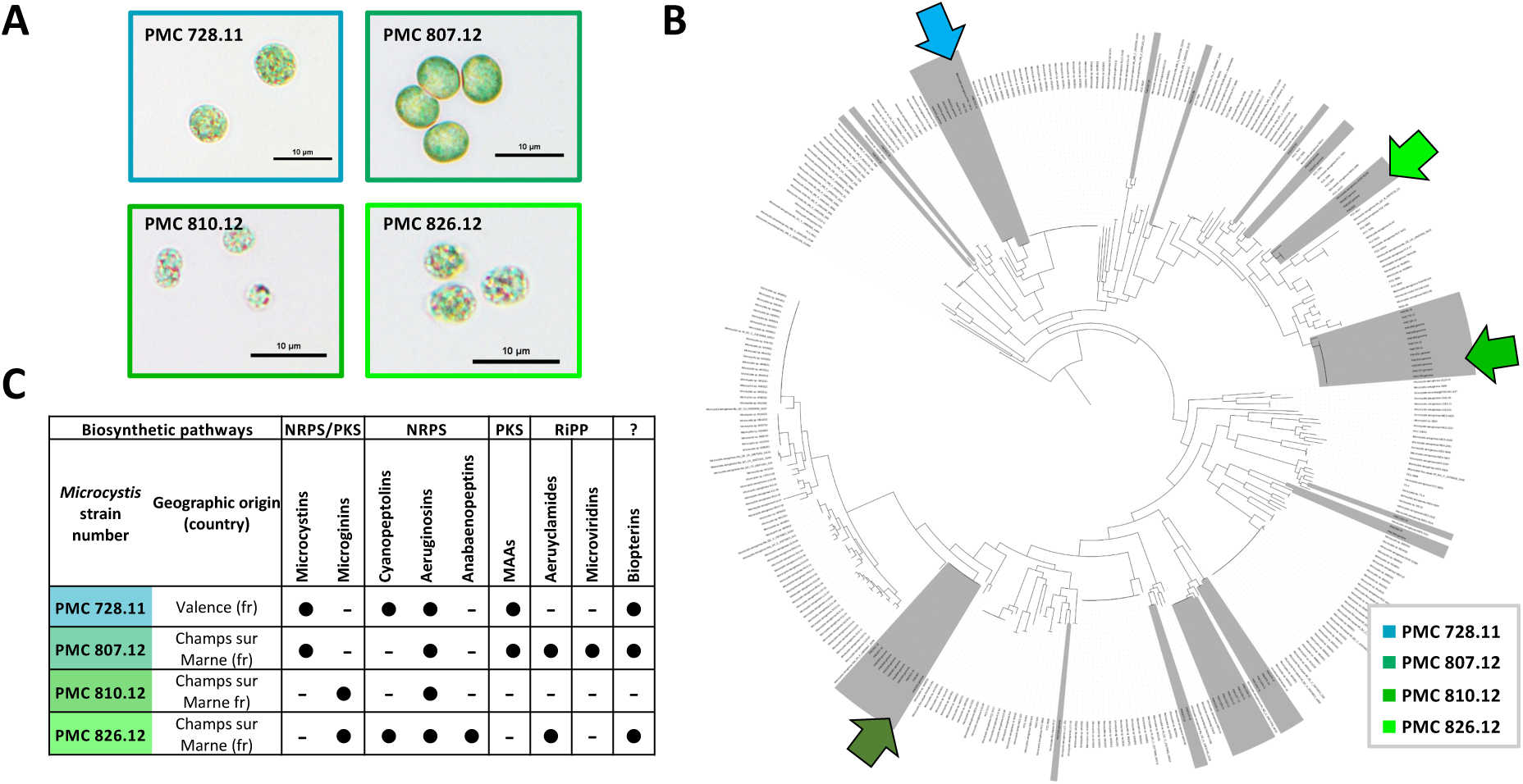
Diversity of the four *Microcystis* strains selected for the present study, regarding their respective morphology under light microscope :(A) their phylogenetic position within main genetic clades regarding the whole phylogenomy of known *Microcystis* genomes, (B) their chemical composition according to direct detection of metabolite chemical families in 75% methanol extracts by mass spectrometry (supplementary figure S1), that were further corroborate with BGC genome contents explored with AntiSmash 6.0 (C). The PMC *Microcystis* strains collected from various French lake are highlighted in grey. Dark spots indicate the production of the different accessory metabolite families.

### Toxicity on Medaka fish embryos and larvae

We performed two different bioassays on Medaka fish embryos (with fertilized eggs from 0 to 2 dpf) and larvae (with hatching larvae from ∼8 to 9 dpf) to document two different toxicological endpoints considering different aspect of the early fish development and the potential associated toxicological impairs (Figure 2). Firstly, the embryo assays performed from 0 to 2 dpf offer the possibility to observe, at the individual level, toxic effects on first developmental stages (considering both embryo death and developmental arrest). This first test showed important effects of the extract of the PMC 810.12 strain on embryo death or developmental arrest, when other strains showed only faint (PMC 728.11 and PMC 807.12) or even no significant toxicity of the PMC 826.12 (compared to control performed with solvent). Regarding teratogenic impairs, only the extract of the PMC 826.12 strain exhibited important teratogenic effects on embryos, observed on tail torsion or head mis-absence (100% of the observed embryos, when other strain extracts didn’t show similar effects on any embryo (n=40)). Secondly, the larvae toxicity tests led to the observation of rapid toxic effects (consisting in an arrest of heart beat and blood circulation). Interestingly, this test performed on well-developed organisms presenting diverse already functional organs, but that do not possess chorionic barrier anymore, shows quite different toxicological responses. Indeed, this second test highlights the overall toxicity of the PMC 728.11 and PMC 810.12 extracts, with 100% of larvae dying within the first 1-2 hours, whereas PMC 807.12 and PMC 826.12 extracts presented low (25%) and no larvae toxicity (n=24), respectively.

**Figure 2.**
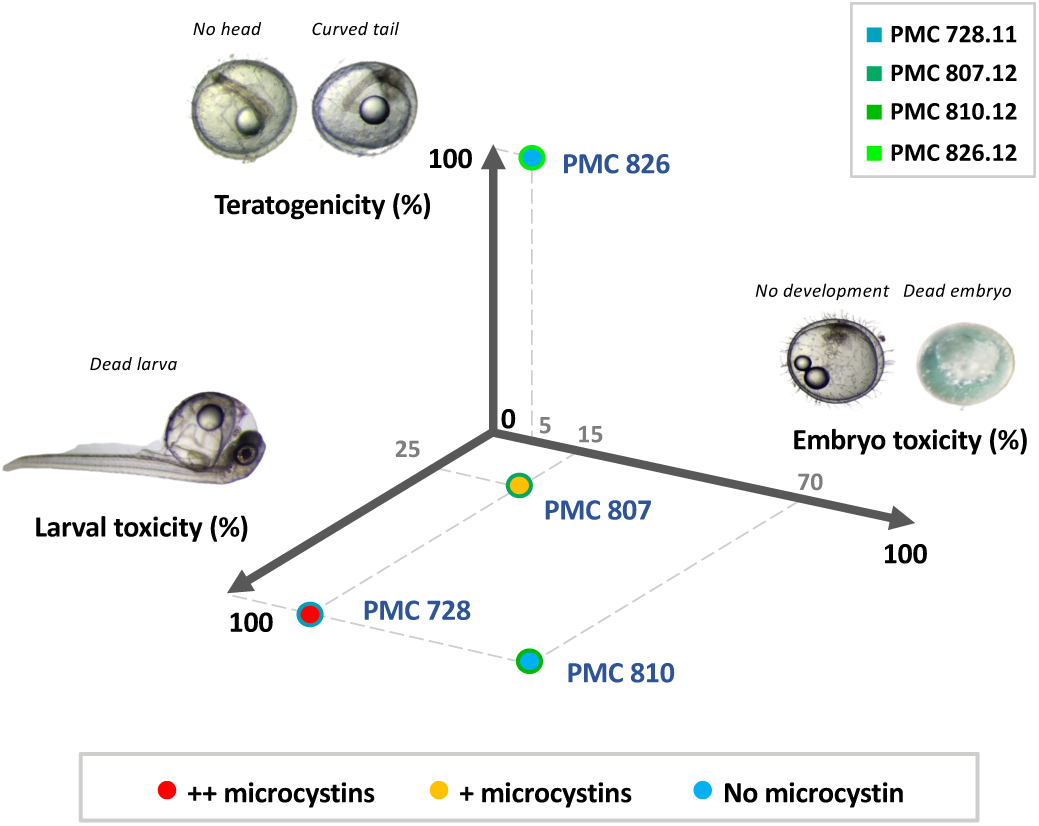
Synthetic results of the toxicity tests performed with the four *Microcystis* extracts selected on the embryos and larvae of the Medaka fish expressed in %age of the embryo death or developmental arrest, the embryo malformation occurrence on tail or head, and the death of larvae. The red, orange and blue colours illustrate the respective MC content rate of each extract.

These apparent discrepancies between the toxicological effects observed with the PMC 728.11 strain extract on embryos (0-2 dpf), on one side, and on the hatched larvae (8-9 dpf), on the other side, could be related with several key aspects of these different tests. Indeed, early embryo toxicity could rather refer to cytotoxic events affecting cell multiplication or survival, whereas larvae present distinct functional organs comprising various differentiated cell types, that could express more diverse potential molecular targets, not expressed in earlier developmental stages. This would then explain higher toxicity observed on larvae presenting more diverse potential molecular targets when compared to the embryos (GopalaKrishnan et al., 2008). An alternative scenario could be proposed when considering the protective effect of the chorion that constitutes an efficient barrier to various contaminants preventing their toxicological effects by precluding their interaction with potential intracellular molecular targets (Mandrell et al., 2012, Yang et al., 2020). Large bio-active cyanobacterial metabolites, such as the heptapeptide MCs, are supposed not to cross the chorion and penetrate to the embryo, thus presenting lower toxicity on chorion-bearing embryos than on post-hatching larvae (Liu et al., 2002). However, in the present case, the semi-quantification of the MC content of the respective strain extracts show that PMC 728.11 extracts contain above 10 times higher MCs levels compared to PMC 807.12, meanwhile the PMC 810.12 and PMC 826.12 extracts present no MC. This observation suggests that certain toxic compounds, other than MCs, might induce the effect observed on embryos exposed to the PMC 810.12 extract, or provoke the teratogenicity observed with the exposure of the 0-2 dpf embryos to the PMC 826.12 extract. Taken together, these results illustrate the remarkable diversity of potential toxicological effects that can be found in different *Microcystis* genotypes, owing to them producing distinct bio-active metabolite corteges (Figure 2).

Several efforts have been performed to develop efficient bioassays on model fish embryos to evaluate cyanobacteria toxicity based on the OCDE 236 fish embryo acute toxicity test (OCDE 2013; Oberemm et al., 1997; Oxendine et al., 2006; Berry et al., 2007; Palikova et al., 2007). Considering the different toxicological effects retrieved form these various fish embryonic tests, one can notice the potential impacts of *Microcystis* on survival, cellular damage, heart beat rate, hatching success, body length, yolk edema occurrence or their behavior, together with the occurrence of various severe developmental perturbations and defects (Saraf et al., 2018; Davidovic et al., 2023; Li et al., 2021). Interestingly, several compounds, such as retinoic acids and carotenoid glycosides, commonly detected in cyanobacteria, including several *Microcystis* strains (Wu et al., 2012; Jonas et al., 2014; Priebojova et al., 2018), have also been shown to induce teratogenic effects on fish early embryonic stages (Jaja-Chimedza et al., 2017). However, the chemical analysis of the molecular diversity of the extracts from four *Microcystis* strains performed herein (supplementary figure S3) does not allow to identify such components, and further effort is required to identify the metabolite that induces the observed teratogenic effects within the PMC 826.12 extracts. So far, the candidate list of *Microcystis* metabolites been potentially the most deleterious according to our fish embryo tests comprises different peptides, including microcystins, cyanopeptolins, microginins or microcyclamides, together with other small compounds such as retinoic acids or di-ter-buthylphenol (Fernandes et al., 2019; Nunes de Freitas et al., 2023; de Almeida Torres et al., 2023; Jonas et al., 2014; Wang et al., 2024). But other *Microcystis*-produced toxic compounds may still have to be characterized.

### Adult Medaka fish exposure and sublethal effects on the microbiome and metabolome

Six-month-old adult female Medaka were exposed four days to live cultures of the four *Microcystis* strains at a standardized concentration of 100 µg.L^-1^ of chl *a*, that is representative of bloom proliferation conditions frequently encounter in temperate aquatic environments during the summer season (Marie and Gallet, 2022). These conditions were initially chosen according to our previous experimentations performed with PMC 728.11 strain (Gallet et al., 2023), with the objective to induce observable, but also ecologically relevant (Le Manach et al., 2018), molecular alterations on at least both gut microbiome and metabolome. We also focused on early effects by favoring short, but sufficiently long, exposure duration (Foucault et al., 2021), in order to minimize acclimatation or indirect side effects, and to observe the molecular response of early stress markers (Gallet et al., 2023). We thus attempt to finely compare the respective specific responses provoked by the different *Microcystis* strains at the level of individual fishes (Duperron et al., 2023). Although this ecotoxicologically relevant experiment doesn’t induce any fish death nor noticeable perturbation of their apparent physiological and behavioral condition, clear impact on both microbiome composition and metabolome content was detected and further examined (Figure 3-6; supplementary figure S4-S14; supplementary table S5-8).

**Figure 3.**
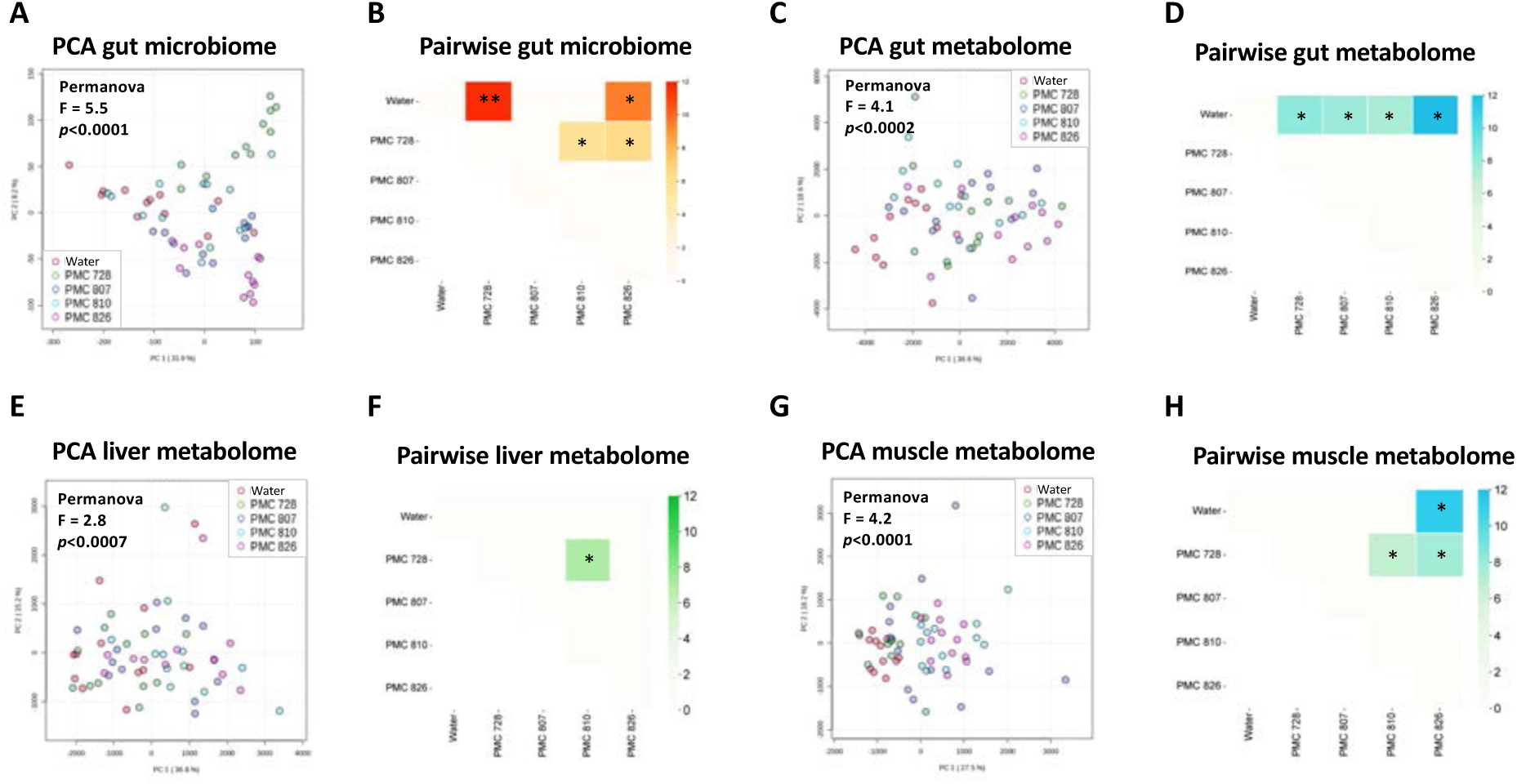
Un-supervised multivariate analyses of the gut microbiome taxonomic composition (A-B), the gut metabolome composition (C-D), the liver metabolome composition (E-F), and the muscle metabolome composition (G-H), regarding individual plots of principal component analyses (PCA), Permanova (A, C, E and G) and pairwise Pairwise Adonis analyses (B, D, F and H). * and ** indicates p-values < 0.01 and 0.001, respectively.

Characterization of the bacterial diversity of the gut microbiome by high-throughput sequencing of the V3-V4 region of the 16S rRNA encoding gene yielded a total of 1,223,933 paired-end quality filtered reads (average 20,398 reads per sample, supplementary figure S4). Species richness appears higher in the gut microbiomes of *Microcystis* exposed fishes compared to those of the water exposed group, and even particularly higher in fish exposed to the PMC 826.12 strain (Supplementary figure S5A). Main bacterial genus-level taxa comprise *Cetobacterium*, *Gemmobacter*, *Hyphomicrobium*, *ZOR0006* (Firmicutes), *Aeromonas*, *Rhodococcus*, *Microbacterium*, *Legionella* and *Terrimicrobium*, all present among the most abundant genera observed in the microbiome of fish guts from all experimental groups (Supplementary figure S5B-C). Un-supervised PCoA performed with Brays-Curtis distance (Supplementary figure S6) and PCA performed on sum scale scaling data (Figure 3A) revealed distinct clusters of taxonomic bacterial composition. The gut microbiome composition of fishes exposed to the culture of *Microcystis* strains (Permanova p<0.0001), in particular the PMC 728.11 and PMC 826.12 strains, showed highly significant differences when compared to those of the water control group as indicated by Pairewise Adonis investigations (p<0.01 and p<0.001, respectively) (Figure 3B).

LC-MS-based metabolomic investigations performed on the fish guts, livers and muscles allow the detection by MS/MS+ of 1,268, 1,389 and 1,994 analytes, 204, 149 and 126 of which, respectively, having been annotated by GNPS supported MS/MS molecular network analysis as potential lipo-phospho-cholines (LPCs), phosphatidyl-ethanolamines (PEs), amino acids, nucleic acids, bile acids, polysaccharides or glutathione-derivates (Supplementary figure S7). Heatmap representations performed with these specific annotated metabolite datasets show that 57, 20 and 50 specific metabolites present significant differences (ANOVA p<0.01) within the 5 experimental groups, in guts, livers and muscles, respectively (Supplementary figure S8). The hierarchical classification trees illustrate the specificity of metabolomes of fish exposed to *Microcystis* strains, and especially those exposed to PMC 826.12 that present the most distinct gut and muscle metabolomes. Similarly, non-supervised multivariate analyses, such as PCA, highlight that the metabolome of certain groups significantly differ from other ones (Figure 3C-H). This is especially the case of gut metabolomes for which the four *Microcystis* strains exposed groups appear significantly different from those of the water control group (Permanova, p<0.001 and Pairwise Adonis p<0.01, Figure 3D). Similarly, the muscle metabolomes of the fish exposed to the PMC 826.12 strain significantly differ from those of the water and the PMC 728.11 groups (Pairewise Adonis p<0.01), whereas the muscle metabolome of fish exposed to the PMC 810.12 strains also diverges from the water control group (Pairewise Adonis p<0.01, figure 3H). For the liver metabolomes, those of the fish exposed to PMC 728.11 strain are the only one presenting significant difference with the PMC 810.12 group (Pairwise Adonis p<0.01, Figure 3F). Regarding the importance of metabolome distinctions between the experimental treatments, one can notice that gut and muscle present highest Permanova F-values (4.1 and 4.2, respectively), whereas liver F-values is only around 2.8 suggesting less marked differences induced by the experimental treatments on this organ. Remarkably, gut microbiome composition presents even more pronounced differences (F-value= 5.5). Taken together, these observations show that the strains inducing the most pronounced metabolic effects are PMC 826.12 and PMC 728.11. Interestingly, PCA projections and hierarchical classifications both indicate that the effect induced by these strains should be largely distinct and rather strain-specific (Figure 3).

The microbial and metabolite variations induced by the exposure to the different *Microcystis* strains on female adult Medaka at both the microbiome and metabolome levels highlight the strain-specificity of these effects, and additional differential analyses were performed to gain further insides into this response specificity at the levels of ASVs or genera and of metabolites (Figure 4-5, supplementary figure S9). First, LEfSe analyses performed at the genus level highlighted the gut microbiome taxa being over-or under-represented among the 5 different experimental groups (Figure 4A). The most dys-regulated bacterial taxon is *Cetobacterium*, whose abundance drops in the gut of fishes exposed to all four *Microcystis* strains (Figure 4B). This genus is generally associated with healthy fish microbiota and may notably contribute as a B_12_ vitamin producer (Tsuchiya et al., 2008; Ma et al., 2019; Duval et al., 2022). *Cetobacterium* is most likely a fish gut resident (supplementary figure S10-S11) and has previously been reported to be stable in abundance in Medaka fish gut upon exposure to limited amount of pure microcystin-LR or cell extract of the *Microcystis* strain (Duperron et al., 2019). Thus, its decrease with fish exposed to the *Microcystus* cultures could be related to potentially dysbiotic mechanisms. Differently, *Gemmobacter*, another abundant bacterium in fish gut microbiotes (supplementary figure S11) (Sheu et al., 2013; Siriyappagouder et al., 2018), exhibits increased abundance when fish are exposed to the four *Microcystis* strains, especially in the PMC 728.11 exposed group (Figure 4B). *Gemmobacter* has previously been observed to increase its abundance under *Microcystis* PMC 728.11 exposure and was, interestingly, co-increasing with other bacteria including *Aeromonas* and *Legionella* (Gallet et al., 2023), as is also observed here (Figure 4B). These latter opportunistic bacteria are often described as potential pathogens (de Bruijn et al., 2018) and their increase could thus be related to the occurrence of dysbiosis situations. A similar increase in opportunistic bacteria was recently documented in guts of zebrafish exposed 96h to *M. aeruginosa* (Qian et al., 2019). In parallel, *ZOR0006* (Firmicutes) also present peculiar abundance variations induced by *Microcystis* strains exposure. Indeed, this fish gut specialist bacterium, that may contribute to the microbiote metabolism by providing specific carbohydrate degradation capabilities, was previously and repeatedly described to decrease upon exposure to PMC 728.11 strain (Foucault et al., 2022; Gallet et al., 2023). In the present case, *ZOR0006* noticeably decrease when exposed to both PMC 728.11 and PMC 826.12, but on the other hand it also increased sharply in guts of fish exposed to PMC 807.12, suggesting that this latter strain may somehow favor the development of this specific bacteria. Finally, unexpected over-abundance of certain bacteria, namely *Hyphomicrobium* and *Microcystis*, in the gut of fish exposed to the PMC 826.12 strain was oberved (Figure 4). Although, it is not entirely surprising to find *Microcystis* ASVs in the gut of fish that have been placed in balneation with *Microcystis* cultures, it is more unexpected that it is only the case with the PMC 826.12 strain and not any of the other three. Such absence of *Microcystis* ASVs in the gut microbiota of fish previously exposed to PMC 728.11 had been interpreted as resulting from a full digestion of all *Microcystis* cells transiting through the digestive tract (Gallet et al., 2023; Foucault et al., 2022). This observation would imply that digestion of *Microcystis* PMC 826.12 cells would not have occurred (this latter assumption been further discussed above regarding other results). Complimentary, PLS-DA analyses have been performed to highlight features contributing the most to the separation of the experimental groups regarding the gut microbiome, together with the metabolomes of the guts, livers and muscles (Figure 5). This differential analysis confirms at the level of the ASVs the observation performed at the genera level with PCoA or PCA (Figure 3A, supplementary figure S6). It also confirms the results of the LEfSe analysis by presenting, as it overall showed various ASV corresponding to *Cetobacterium*, been more abundant with the water control treatment, and many other corresponding to *Gemmobacter*, *Aeromonas*, *Terrimicrobium* and *Pirellula*, among others, been over-represented in fish guts from the PMC 728.11 strain treatment (Figure 5B). Regarding the metabolomes of guts, livers and muscles, the PLS-DA individual plots overall depict a greater distinction of the metabolomes of fishes exposed to the PMC 826.12, which appear more isolated from the metabolomes of those exposed to control condition than the three other *Microcystis* strain groups (Figure 5C, E and G). Firstly, concerning the guts, metabolites that explain most of the group distinction (best scores of variable importance in the PLS-DA projection, VIPs) correspond to various LPCs, PEs, cholic acids and peptides, that present higher abundances in fish exposed to either water control or PMC 826.12 groups (Figure 5D). Secondly, regarding the livers, metabolites most involved in the group distinction correspond principally to LPC, PE and nucleic acids, that present higher contents in fish exposed to PMC 810.12 and PMC 826.12 (Figure 5F). Thirdly, about the muscles, metabolites most involved in the group distinction mainly correspond to nucleic acids and saccharides, potentially related to various metabolic processes occurring in fish exposed to the four different *Microcystis* strains (Figure 5H) being potentially related with detoxification events.

**Figure 4.**
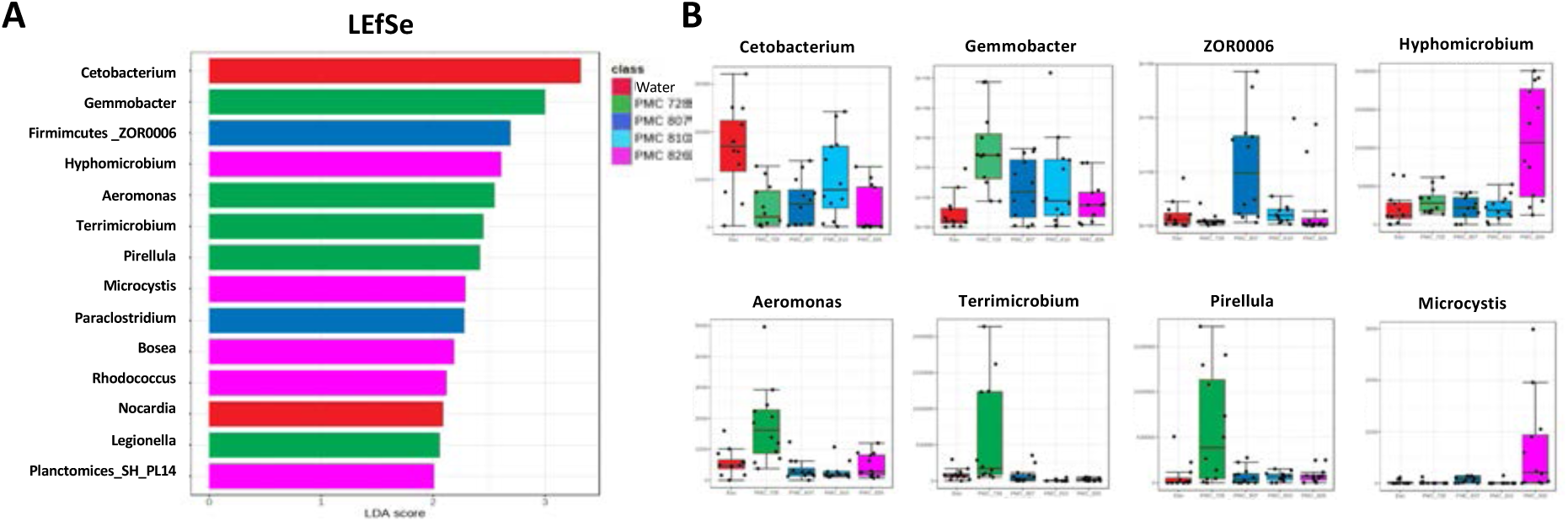
Effects of the exposure of the young female adult Medaka fish exposed 4-days to the different *Microcystis* strains cultured to the taxonomic composition (performed here at the family level) of the gut microbiome according to comparison to control fish exposed to water on barplot of LEfSe discriminant analysis (p<0.01 and LDA score>2) (A) and respective boxplots of the 8 most dys-regulated genera (B).

**Figure 5.**
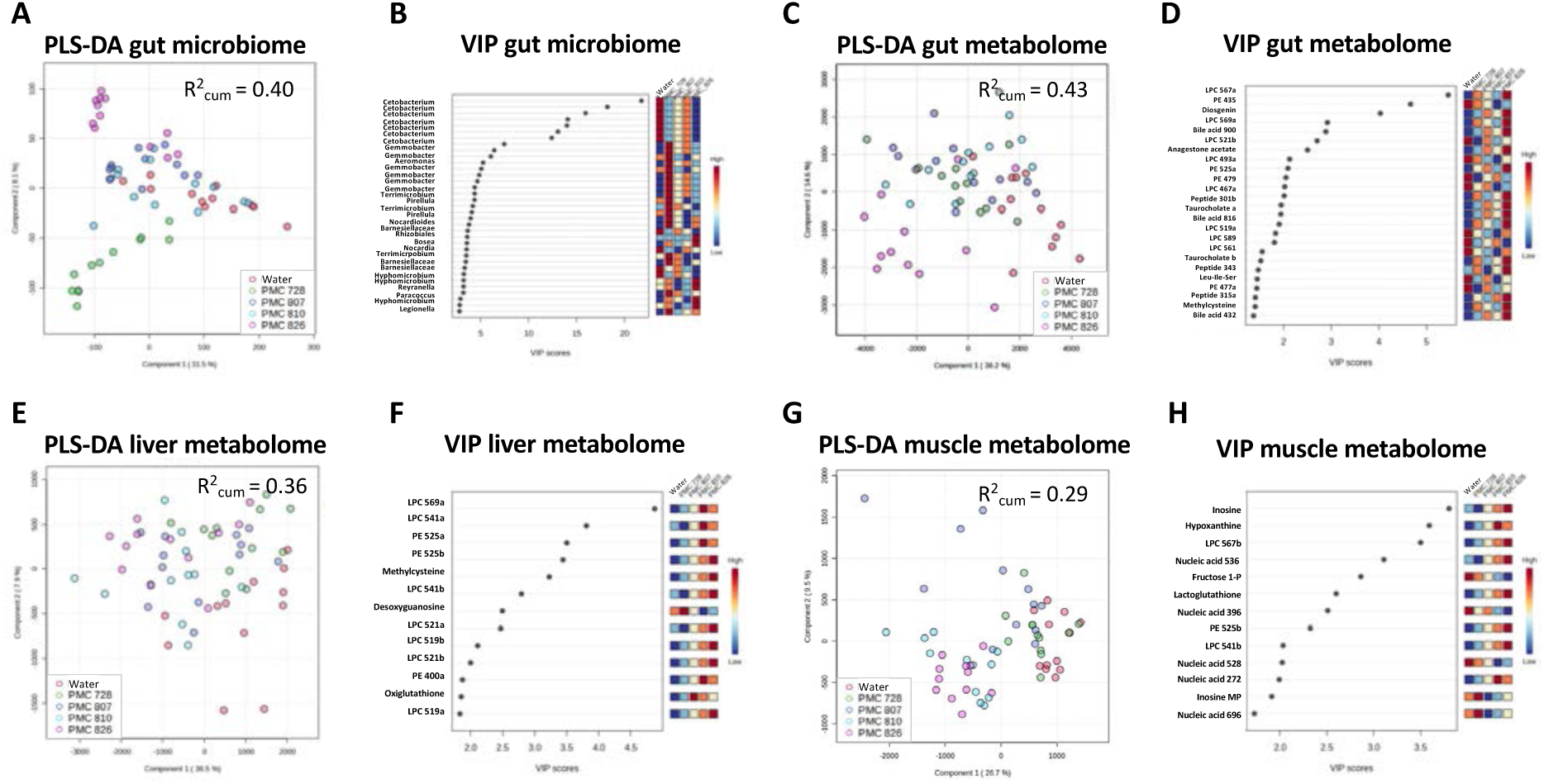
Supervised multivariate analyses performed on the composition of the gut microbiome (A-B), the gut metabolome (C-D), the liver metabolome (E-F), and the muscle metabolome (G-H), regarding individual plots of partial least-square differential analysis (PLS-DA) (A, C, E and G) and highest ranked scores of the respective variable importance in projection (VIP) (B, D, F and H).

### Multi-omic integration of the fish molecular responsiveness to *Microcystis* exposure

On one hand, considering the whole dataset (including gut microbiome together with gut, liver and muscle metabolome of fish exposed to the four different *Microcystis* strain and the water control) provides a general correlation scheme (Supplementary figure S12) and depicts the global connections between the different features within the organism responding to the different stresses induced by the exposure (Supplementary figure S13). This approach exquisitely illustrates the co-variation correspondanec between different metabolites and bacteria, although it does not allow to determine the causality nature of the relation between these linked features. Overall, it distinctly shows: 1/ negative correlations bewteen 5 *Cetobacterium* ASVs and 9 cholic acids in the guts; 2/ negative correlations between the variation of 7 *Gemmobacter* ASVs abundances and various lipids in the guts (including LPCs and PE); 3/ positive correlations between *Microcystis* and *Hyphomicrobium* ASVs with 20 different peptides from the guts.

On the other hand, the experimental conditions considered separately (*e.g.* the four different *Microcystis* strains compared to the water control) rather highlight the specific covariations resulting from the specific response to the different strains (Figure 6A-D). Firstly, exposure to PMC 728.11 strain is associated with numerous negative co-variations between abundances of *Cetobacter* ASVs and several LPCs and cholic acids from guts, and several positive co-variations of *Gemmobacter* ASVs with various LPCs, PEs and peptides (Figure 6A). Interestingly, certain metabolites from muscles (naphtalene-dihydrotiol and Hexamethoxyflavone) exhibits pivot positions characterized by a large set of numerous positive or negative correlations with various other features from other organs and with different bacteria from the gut microbiota. Secondly, the fishes exposed to the PMC 807.12 show a less complex pattern comprising lesser features or correlation links (Figure 6B). Thirdly, fishes exposed to PMC 810.12 exhibit positive correlation between 7 *Gemmobacter* ASVs, 3 LPCs and 8 cholic acids from guts, together with 10 LPCs and 3 PEs from the livers (Figure 6C). At last, the fishes exposed to PMC 826.12 strain present a clearly distinct correlative pattern that comprises 7 *Cetobacterium* ASVs of which abundances are negatively correlated with those of 19 peptides and 6 cholic acids from the guts, whereas *Hyphomicrobium* ASVs show positive correlations with the same 19 amino acids and 6 cholic acids from the guts, plus 6 LPCs and 2 PEs from the livers, together with positive and negative correlations with few amino acids, saccharides and nucleic acids from muscles, and with other gut bacteria including *Bosea*, *Pseudonocardia* and *Rhizobiales* among others ASVs (Figure 6D).

**Figure 6.**
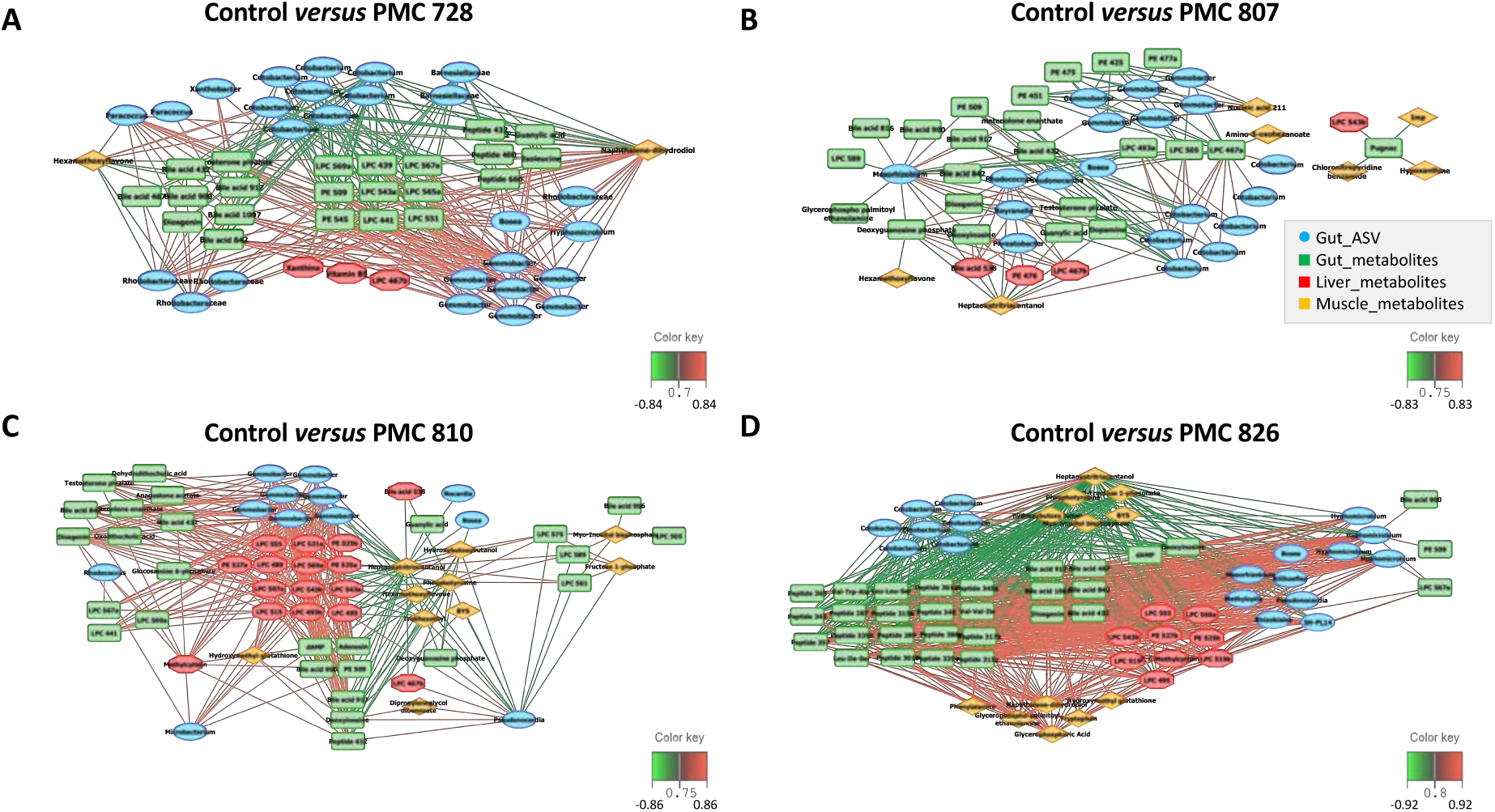
Relevance network analyses performed with MixOmics illustrating the most correlated metabolites (green, red, and orange for gut, liver and muscle metabolome) and gut ASVs (in blue) discriminating among the 4 treatments compared to the control water treatment considered separately (A for PMC 728.11; B for PMC 807.12; C for PMC 810.12 and D for PMC 826.12). Only variables (ASVs and metabolites) with correlation values above ±0.77 are displayed. Pearson correlations between covariate metabolites and ASVs are represented by coloured segments (green: negative association, red: positive association).

Precision ecotoxicology approach integrating various biological compartment at the individual levels offers unprecedent perspectives for understanding the fine response to various environmental stressors (Brooks et al., 2023). In the present case, it allows to compare the specific effects of different *Microcystis* strains on both physiological impairs and gut microbiota dysbiosis (Marie, 2020). Also, a microbiome-aware ecotoxicology taking into account the holobiont dimension on macro-organisms would benefit from better considering the microbial dynamic interface between the organism and its environment (Duperron et al., 2020). The gut microbial community disruption is associated with potential metabolic changes, that are observed at the level of gut metabolome. These variations indicate the occurrence of dysbiosis outcomes, with potential physiological consequences for the holobiont that can be subsequently perceived on other fish organs. In the present case, although the liver metabolome displays limited signs of perturbation, very likely because of the short duration of the experiment (i.e. four days of cyanobacteria exposure), the muscle metabolomes show more pronounced variations with remarkable specific molecular signatures.

Our observations overall confirm that the PMC 728.11 strain produces different accessory metabolites that may be involved in the important variations observed in fish gut microbiota (Foucault et al., 2022; Gallet et al., 2023). Basically, the PMC 728.11 strains produce above 10 times more MCs than the PMC 807.12, that could explain some the distinct effects observed. Among those metabolites also already identified, aerucyclamides (*syn.* microcyclamides) are assumed to present potent antimicrobial bioactivities (Martins and Vasconcelos, 2015; Portmann et al., 2008), and could impact the microbiota of the intestine lumen by direct action on certain sensitive bacterial taxa, as previously suggested (Gallet et al., 2023). Differently, the PMC 826.12 strain presents quite distinct impairs as the remarkable dys-regulations observed at both gut microbiome and metabolome levels. It is characterized by a *Cetobacterium* decrease and an outstanding increase of *Hyphomicrobium* and *Microcystis* associated with a large increase of various peptides and cholic acids in the gut, of several LPCs and PEs in the liver and of various small metabolites in the muscle. Taken together, these observations highlight the broad and fundamental differences in the molecular responses occurring in different organs of fish exposed to the different *Microcystis* strains, staring both microbiome and metabolome.

### Digestion inhibition induced by the *Microcystis* PMC 826.12 strain?

The LC-MS-based investigations of faeces metabolite contents show the presence of several molecules specific to *Microcystis* strains cultures, including aeruginosins, aerucyclamides, microginins, anabaenopeptins and cyanopeptolins (Supplementary figure S14A). Comparison of the faeces metabolomes also reveals the remarkably higher metabolite content in the faeces of fishes exposed to the PMC 826.12, in particular (Supplementary figure S14B). Discriminant analysis indicates that these differences are mostly due to the presence of various *Microcystis* specific metabolites, such as microginins, anabaenopeptins or aerucyclamides present in the PMC 826.12 biomass (Supplementary figure S14C). We notice that the faeces of fishes exposed to the other strains do not similar high *Microcystis* metabolite contents. Regarding the aspect of these faeces, we observed they contain high amount of greenish material, that were also presented in the guts of those fishes (Supplementary figure S14E-F). We thus assume it corresponds to undigested *Microcystis* cells, this hypothesis being further supported by the high abundance of *Microcystis* sequences in the gut microbiota (Supplementary figure S15). Overall, these observations suggest poorly efficient and incomplete digestion of PMC 826.12 *Microcystis* cells, whereas *Microcystis* cells of other stains appear almost fully degraded by the digestive process (Supplementary figure S15).

Interestingly, previous investigations have already highlighted that certain *Microcystis* proliferations could be responsible for digestion inhibition in the ciliate *Blepharisma americanum* (Chapman et al., 2019) or *Daphnia magna* (Agrawal et al., 2005), or negatively impact fish appetite through bioactive components that remains to be identified (Niu et al., 2024). These latter authors proposed that the mitigation effect could be explained by the alteration of intestine amino acid metabolites, that lead to decreasing appetite through bile acid and lipid metabolism regulation. So far, *Microcystis* bioactivity screening has also revealed that several families of their specialized metabolites present *in vitro* digestive trypsin and chymotrypsin protease inhibition (Adiv and Carmeli, 2013). Indeed, several variants of both aeruginosins, cyanopeptolins (*syn.* Micropeptins), anabaenopeptins, microviridins and microginins have been shown to exhibit various protease inhibition activities against trypsin-type proteases, principally (reviewed in Demay et al., 2019). Thus, the different cyanopeptolin, aeruginosin, microginin and anabaenopeptin variants produced by the PMC 826.12 strains could be considered as potential perturbators of Medaka fish digestion - the exact molecules remaining to be identified. However, the alternative hypothesis of the induction by *Microcystis* metabolites of primary gut microbiome dysbiosis with cascade consequences on digestion regulation and collateral perturbation on amino acid, bile acid and lipid metabolism also remains plausible and subsequent causality relations remain to be investigated.

### Ecotoxicological outcomes for natural *Microcystis* blooms

By comparing the deleterious effects induce by different strains of *Microcystis*, one of the most commonly proliferating harmful cyanobacterium (Harke et al., 2016), we show how the chemical diversity of produced bioactive metabolites may differently affect aquatic organisms experiencing frequent blooms through their life. Indeed, our multi-endpoint approach evaluating acute effects on embryo/larvae and environmental dose effects on adult Medaka fish emphasizes the wide range of the perturbations expressly induced by the different *Microcystis* chemotypes. In particular, we found 1/ remarkable toxicity or teratogenicity on chorion-bearing embryos induced by the non-MC producing PMC 810.12 and PMC 826.12 strain extracts, respectively, 2/ specific toxicity on chorion-free larvae provoked by the MC-producing PMC 728.11 strain extract, whereas 3/ the PMC 728.11 and the PMC 826.12 strains (and PMC 810.12, in a lesser extent) present more marked dys-regulation on the gut microbiome and metabolome compositions. In addition, 4/ the metabolites produced by PMC 826.12 strains seems to induce specific inhibition of the fish gut digestion. Differently, 5/ the PMC 807.12 strain, presenting low MC contents (above 10 times lower than in PMC 728.11), present lower embryo/larvae toxicity and rather favor an increased proportion of *ZOR0006*, a potential commensal-to-mutualistic member of the fish gut microbiome (Gallet et al., 2023).

Overall, these observations illustrate that MC are not the only threatening *Microcystis* metabolite, and that various other toxicants might be responsible for different effects induced by the four different strains presently explored. Although, it is also likely that the toxic effect induced by the PMC 728.11 extract on larvae would be due to its high content in MCs, which are presumed not to cross the chorion barrier protecting the early embryos, the toxic effects induced by the PMC 810.12 strain on both embryos and larvae is probably mediated by a contaminant of another kind, that can pass through the chorion barrier. This latter strain produces a large variety of secondary metabolite families including microginins, cyanopeptolins, aeruginosins, anabeanopeptolin, aerucyclamides and biopterins, together with other unknown metabolites that still have to be described, according to genome mining and metabolomic approaches (supplementary table S1). These toxic compounds could cross the chorion as they are expected to present lower molecular weight and be less hydrophobic, than MCs or cyanopeptolins. Alternatively, it remains be possible that specific active transporter would be present on the embryos surface, but so far this point has not been confirmed so far regarding egg membrane (Singo et al., 2017; Qiao et al., 2019). Similarly, the molecule responsible of the teratogenic on the early embryo effects induced by the PMC 826.12 extract doesn’t seems to belong to identified members the retinoic family (Wu et al., 2012), as no known variant could have been detected so far from the metabolomic analysis.

A step further, long-term chronic investigations performed with environmental concentrations of these different *Microcystis* strains/genotypes would now offer more accurate evaluation on their respective consequences on fish life traits including developmental success, growth rate, behavior and generally ecological performances (Falfushinska et al., 2021; Cai et al., 2022). Considering that natural blooms might be composed of variable proportions of different *Microcystis* genotypes (Briand et al., 2009; Perez-Carrascal et al., 2019) producing different metabolic cocktails (Lifshits and Carmeli, 2012) they may also represent various threats for fishes (Maric et al., 2020).

However, it remains especially difficult to anticipate and to predict the ecological influences of cyanobacterial proliferations in the different fish populations and guilds (de Almeida et al., 2024; Drobac et al., 2021). Indeed, both MC-producing and non-MC-producing *Microcystis* genotypes have been shown to present deleterious, if not toxic effects, in comparable but also different manners (Le Manach et al., 2018; Sotton et al., 2017). This observation now calls for an in-depth consideration of which metabolites may cause ecotoxicological concerns for the different aquatic organisms beyond MC (Janssen 2019).

Also, it still remains puzzling that robust significant correlation could not be retrieved between cyanobacteria proliferation, MC production and fish tissue contamination (Drobac et al., 2016). Indeed, it now seems pertinent to consider that the concomitant production by natural cyanobacterial bloom of bioactive metabolites with potential digestive enzyme inhibitor capacities together with cyanotoxins, such as MCs, may soundly impact the digestion of the cyanobacterial cell in the fish gut lumen and thus influence it adsorption by the epithelia and the subsequent contamination of the whole organism by the cyanotoxins themself.

## Conclusion

Our present work aims at enlightening the diversity of ecotoxicological impairs that might be considered regarding the intrinsic diversity in terms of genotypes of natural *Microcystis* blooms and of associated metabolite diversity that still been progressively revealed.

The ecotoxicological challenges that now emerges would to be capable to apprehend the consequences of 1/ been able to disentangle the complex cocktail effects occurring with the exposure to multi-cyanobacterial bioactive metabolites, and to predict toxicities of those mixtures, 2/ iterative exposures, as successive blooms could induce a cumulative drift to fish physiology, leading to suboptimal states and health impairments, together with 3/ apprehending the various disturbances that may concern the different fish species, and to evaluating their consequence on fish guild and aquatic ecosystem functioning.

## Acknowledgements

This work was supported by the ANR MC-Tox project, grant ANR CE34-SJ 11008-22 of the French Agence Nationale de la Recherche, and the Tox Model project financed by grant ANSES 22-EST-111 of the Agence National de sécurité Sanitaire de l’alimentation, de l’Environnement et du travail. Maïwenn Le Meur PhD is funded by a DIM One-Health 2.0 grant provided by the Ile-de-France region. The Paris Muséum Collection (PMC) is funded by the Muséum National d’Histoire Naturelle. The mass spectrometry analyses were acquired at the Plateau technique de spectrométrie de masse bio-organique, Muséum National d’Histoire Naturelle, Paris, France. We thank Eglantine Soubrand and Pierre Foucault for their technical assistance.

## Author contributions

BM, CD, EL and SD conceived and designed the experiments; CD isolated all new strains of the PMC; MLM, MQ and MB performed the analysis; CD, MLM, MQ and BM treated the data. All authors wrote and reviewed the manuscript.

## Competiting interests

The authors declare no conflict of interest.

## Supplementary figures

**Supplementary figure S1.**
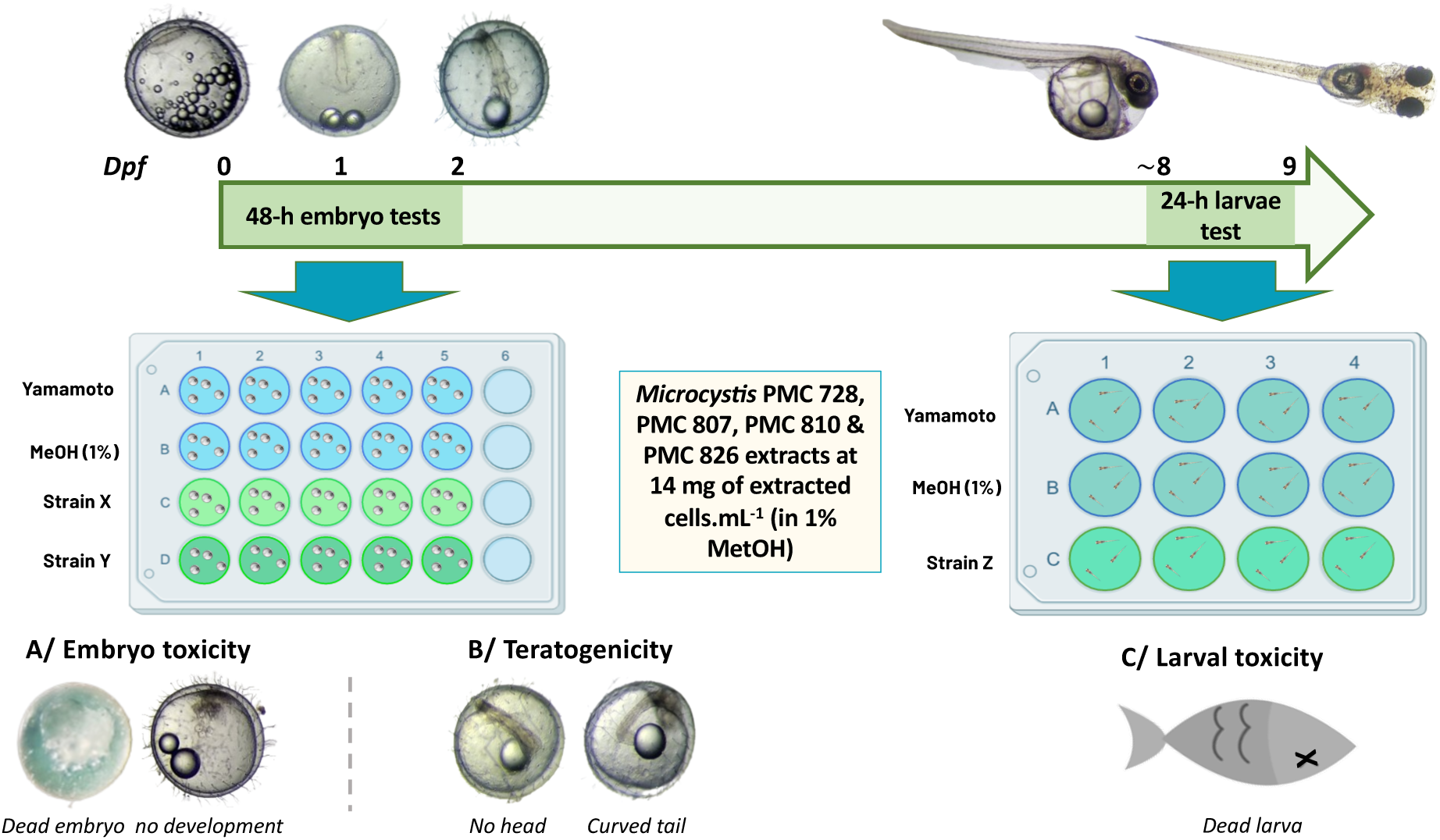
Schematic illustration of the experimental design of the Medaka fish embryos and larvae exposure to the 75% methanol extracts of the 4 *Microcystis* strains selected. All extracts were tested at the same maximal concentration of 14 mg of extracted *Microcystis* cells per milliliter in 1% methanol on freshly fertilized embryos from 0 to 2 dpf and hatched larvae from above 8 to 9 dpf. Toxicity was evaluated on the survival and the malformation rates of embryos (n=4 in 5 replicates) and the death of larvae (n=3 in 4 replicates).

**Supplementary figure S2.**
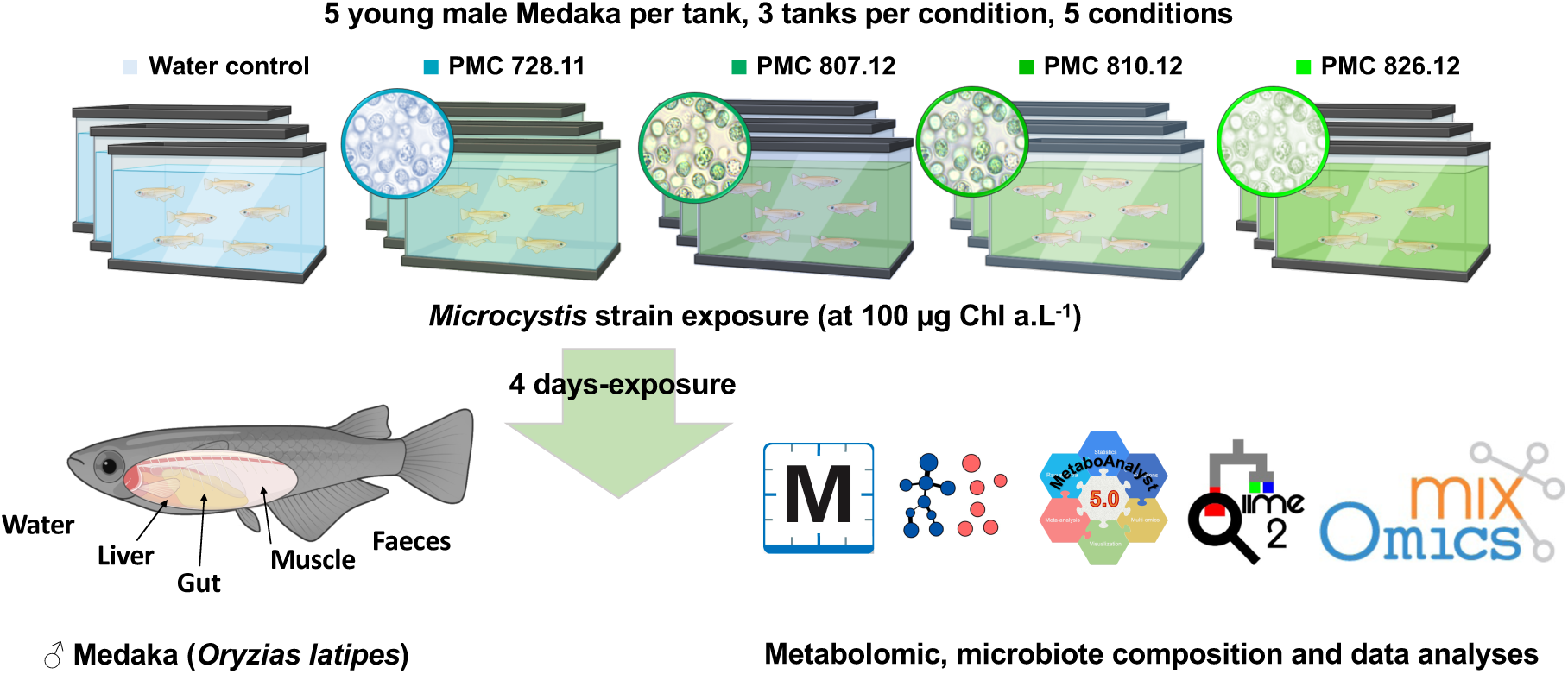
Schematic illustration of the experimental design of the 4-day exposure of the Medaka fish to the culture biomasses of the 4 *Microcystis* strains selected, using freshwater as negative control. All strains were tested in triplicats at the same cellular concentration estimated according the chl *a* concentration of 100 µg.L^-1^ on 5 young female adults per 10-L tank. Water, faeces, gut, liver and muscle samples were collected at the end of the experience for subsequent metabarcoding and metabolomic analyses.

**Supplementary figure S3.**
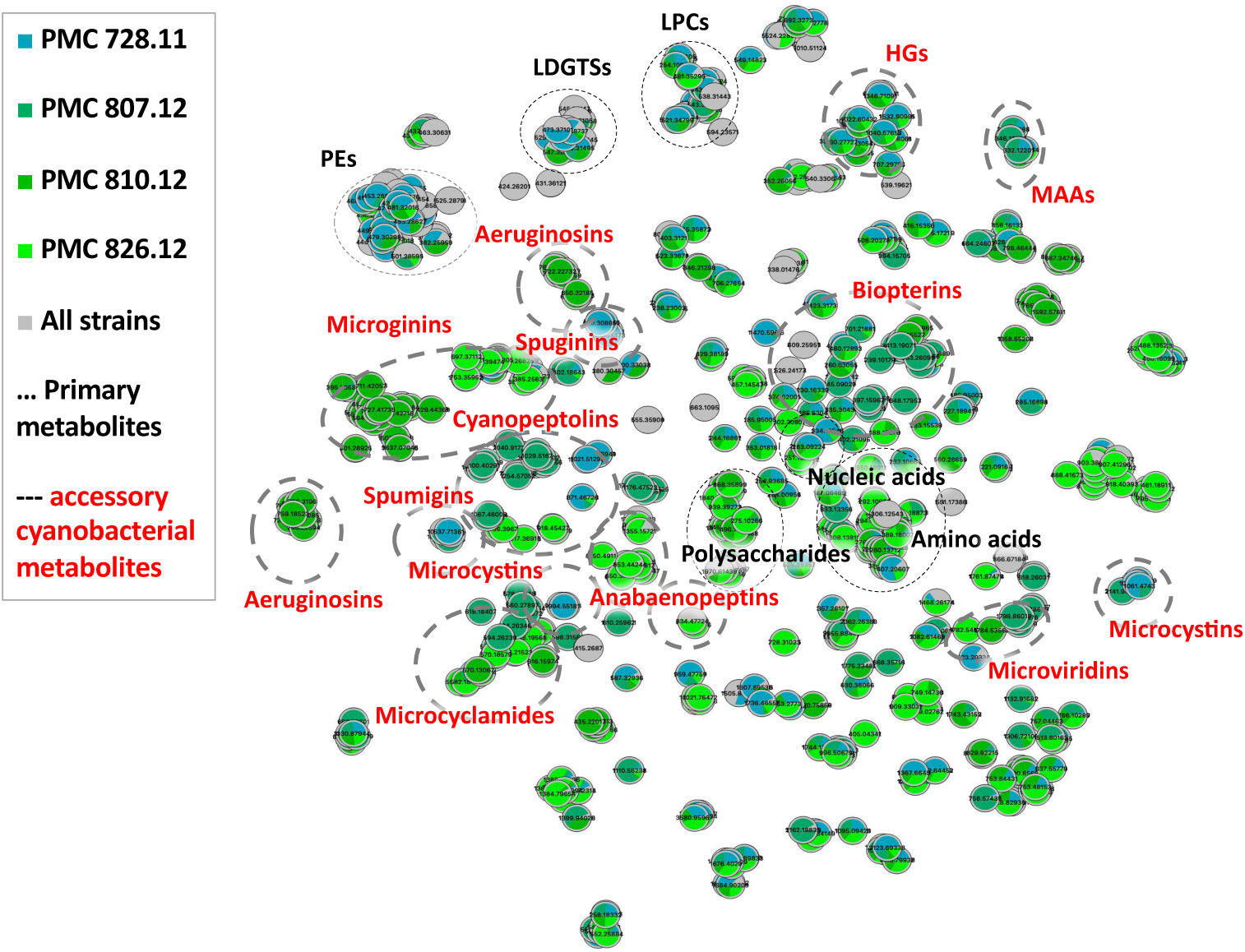
Molecular networking of metabolites extracted from the 4 *Microcystis* culture biomass generated with *t*-SNE algorithm, with specific cyanobacteria and primary metabolite clusters indicated in red and black, respectively (PE, phosphatidylethanolamine; HG, heterocyte glycolipid; LDGTS, lysodiacylglyceryltrimethyl homoserine; LPC, lipophosphocholine; MAA, mycosporine-like amino acid).

**Supplementary figure S4.**
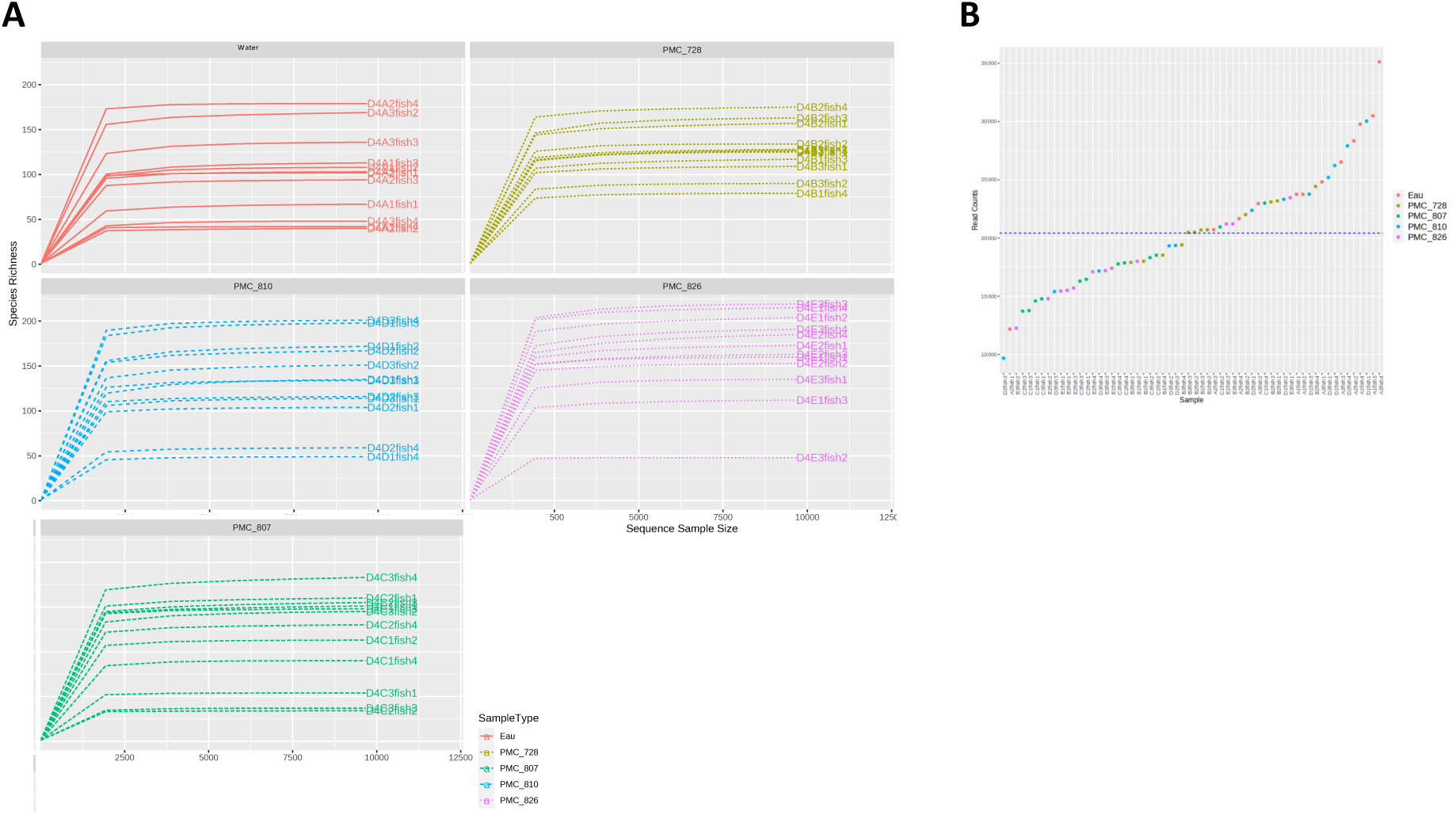
Rarefaction curves of the microbiome gut samples analyzed from the different experimental conditions showing minimum and maximum counts of 9,672 and 35,124 reads, respectively.

**Supplementary figure S5.**
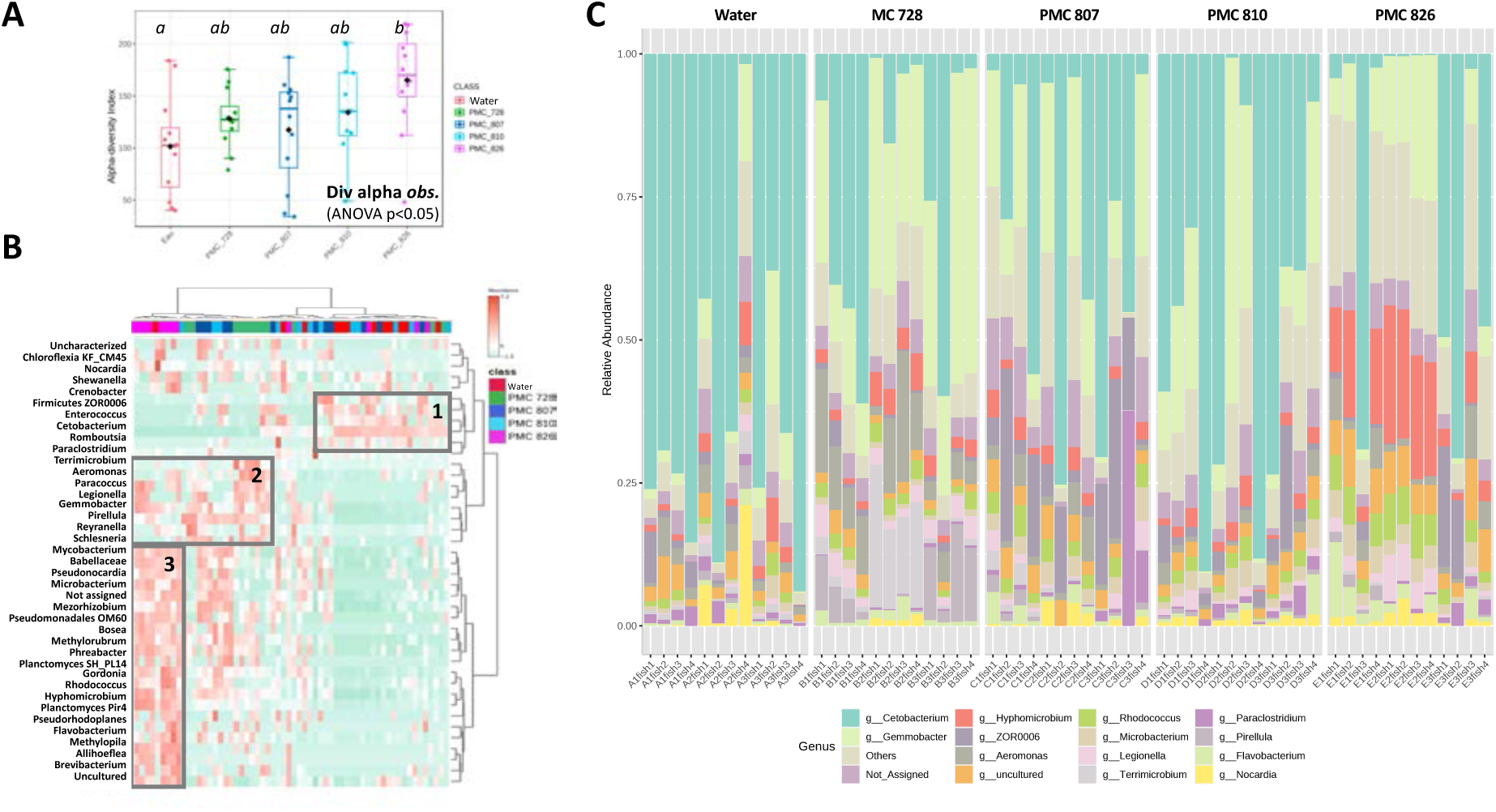
Taxonomic composition of the gut microbiote of fish exposed to the different treatment considering the bacterial genera level shows increasing diversity richness in the gut microbiome of fish exposed to the *Microcystis* strains and in particular with the PMC 826.12 strain (A), relative abondance on heatmap with hierarchical classification considering normalized dataset (B) highlighting taxa over-represented in the gut microbiome of fish rather exposed to control (box1), both the PMC 728.11, PMC 810.12 and PMC 826.12 strains (box2) or the PMC 826.12 strain in particular (box3), together with individual relative composition histogram (C).

**Supplementary figure S6.**
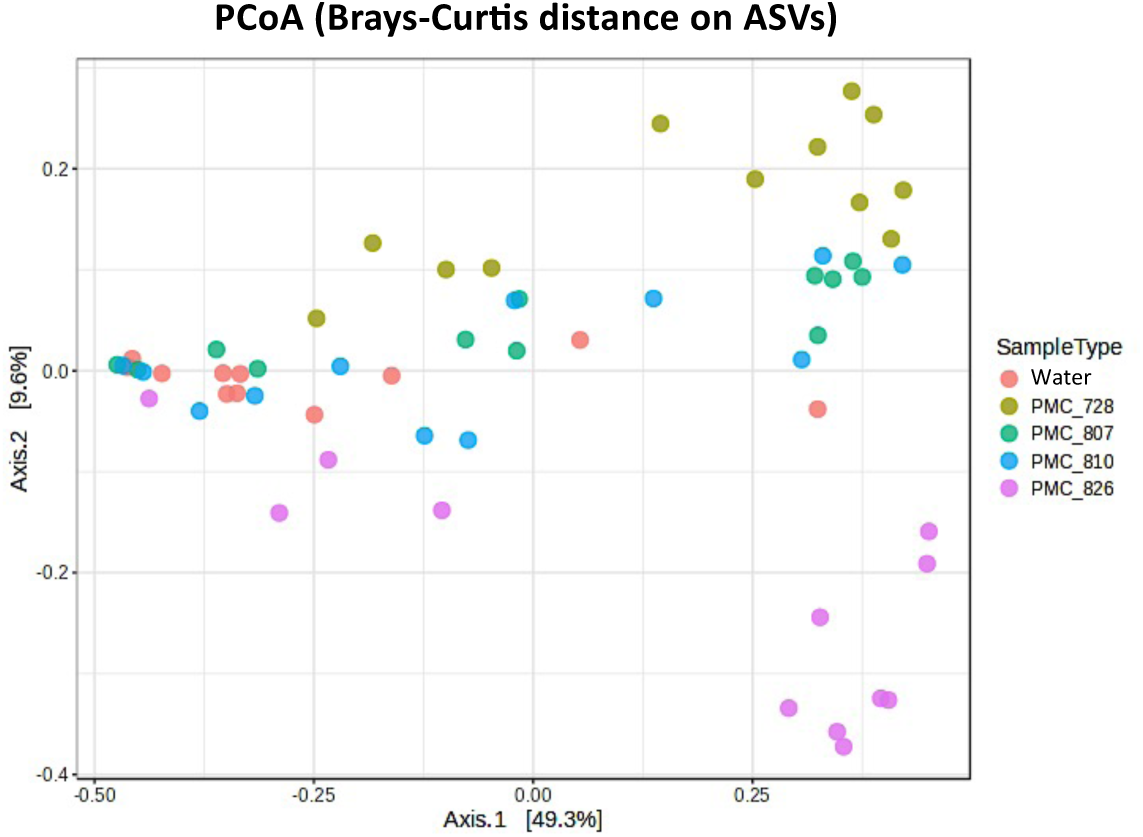
Individual plot of the Principal Coordinate Analysis (PCoA) performed with Brays-Curtis distance showing significant discrimination according to the bacterial composition of gut microbiomes (Permanova p<0.0001; F-value =5.24).

**Supplementary figure S7.**
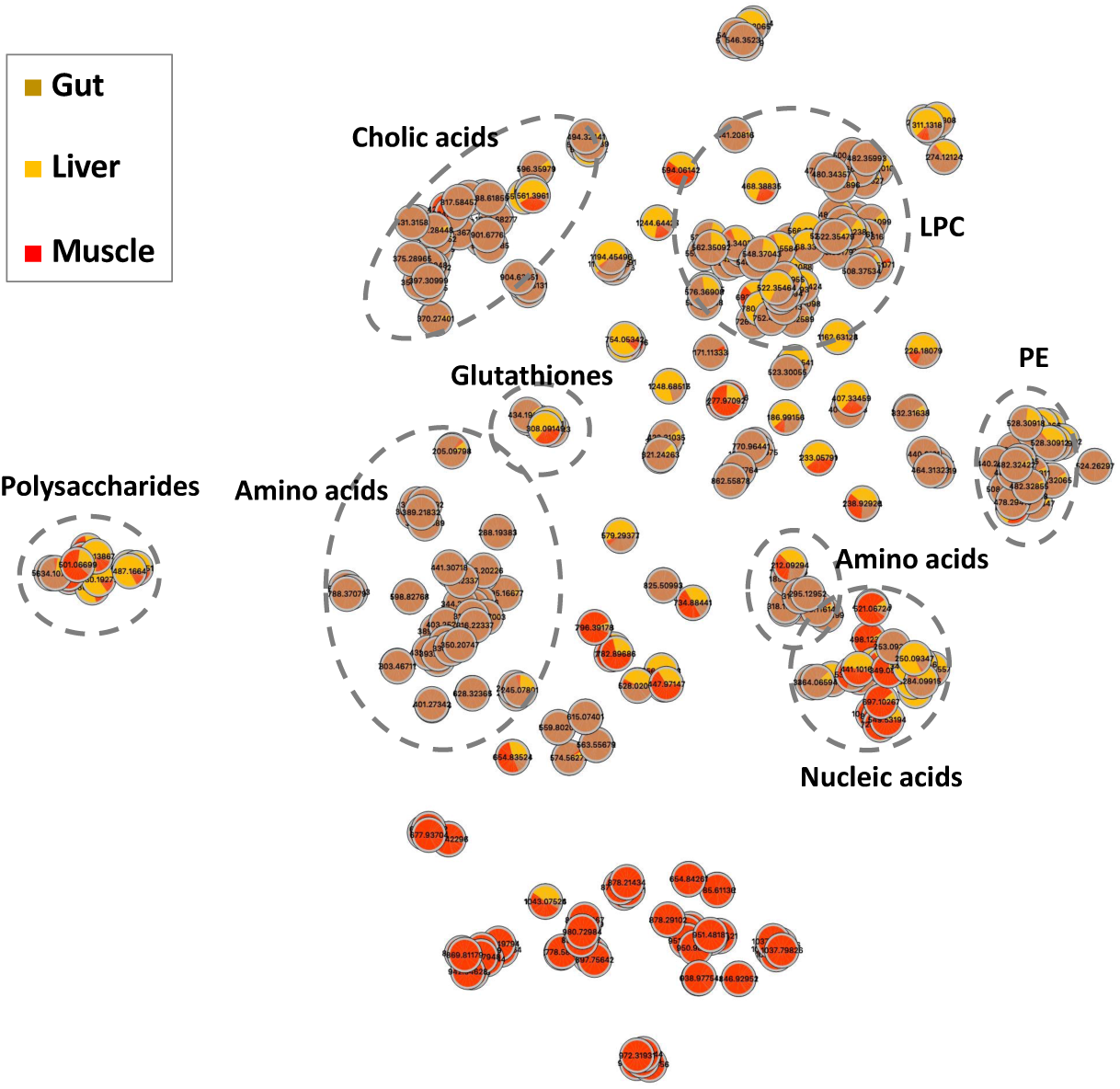
Molecular networking of metabolites extracted from the gut, the liver and the muscle of the medaka fish generated with *t*-SNE algorithm, that supports the annotation of 204, 149 and 126 metabolites, respectively, belonging to cholic acid, lipo-phospho-choline (LPC), phosphatydil-ethanolamine (PE), glutathione derivate, nucleic acid, amino acid or polysaccharide clusters.

**Supplementary figure S8.**
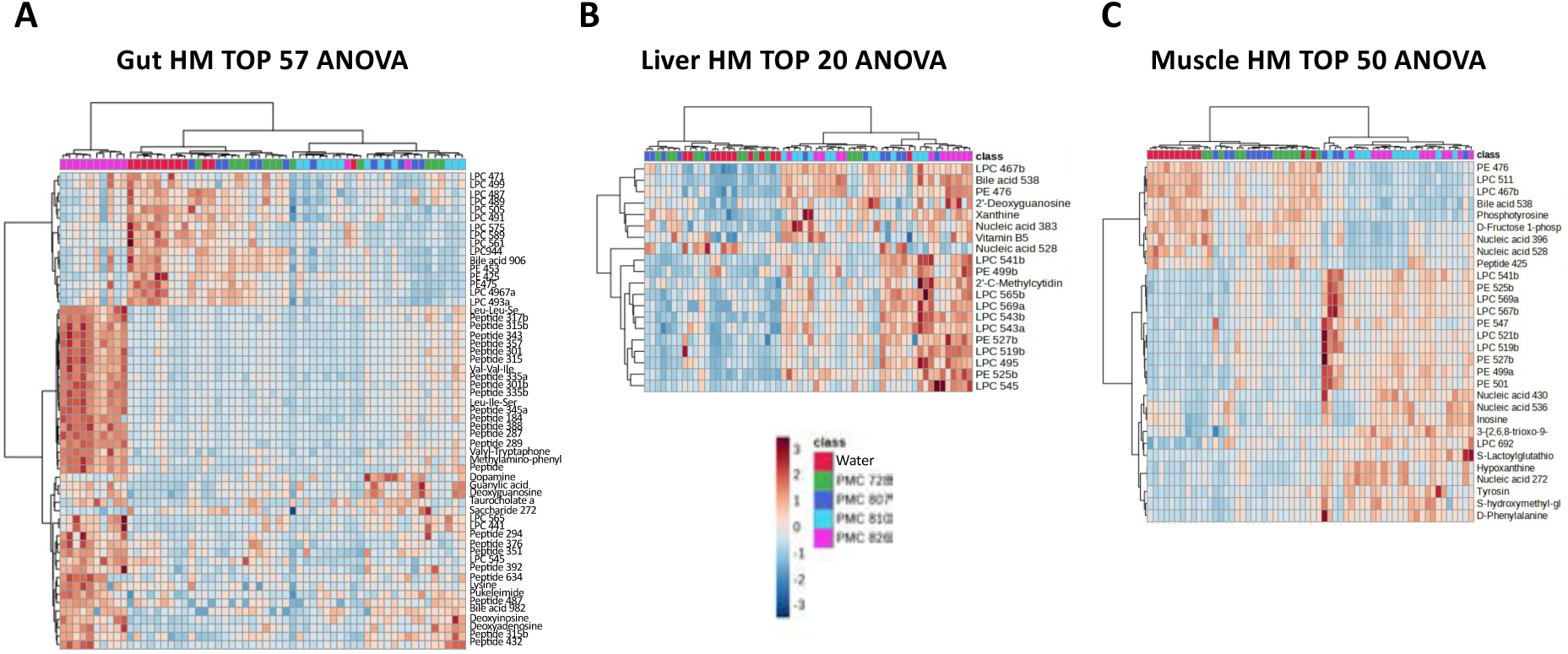
Heatmaps with hierarchical classification of annotated metabolites presenting significantly different distribution (ANOVA p<0.01) in the gut, the liver and the muscle metabolomes of fish exposed 4-days to the different Microcystis strain cultures, representing 28, 13 and 40% of the whole metabolite annotated in the different tissues, respectively.

**Supplementary figure S9.**
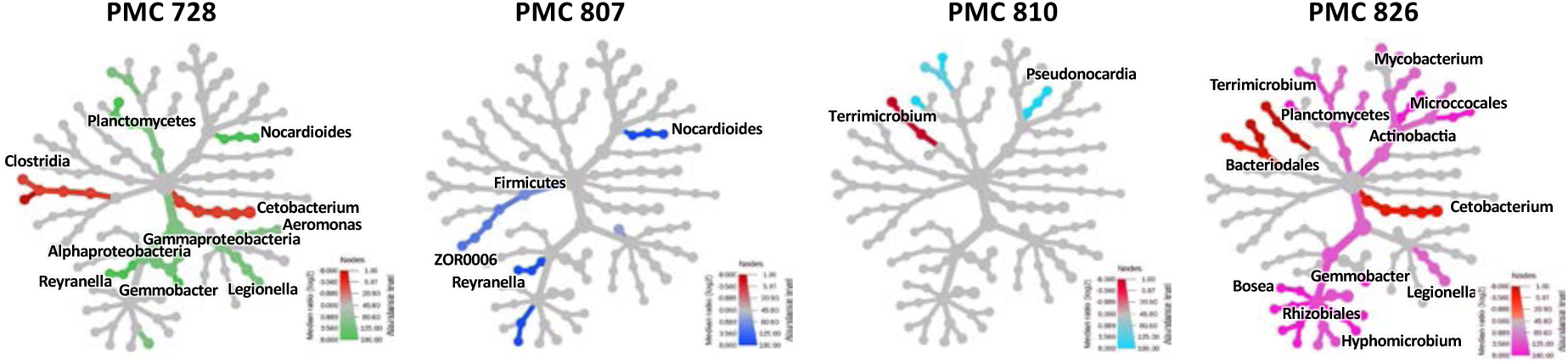
Effects of the exposure of the young female adult Medaka fish exposed 4-days to the different *Microcystis* strains cultured to the taxonomic composition (performed here at the family level) of the gut microbiome according to comparison to control fish exposed to water on respective taxonomic heat trees (non-parametric Wilcoxon test, p<0.05). Green, blue, turquoise, pink and red indicates taxa over-represented in PMC 728.11, PMC 807.12, PMC 810.12, PMC 826.12 and water treatments, respectively. These representations illustrate that both exposures to PMC 728.11 and PMC 826.12 have induce more numerous and specific taxonomic variations on the fish gut microbiome composition than the PMC 807.12 and PMC 810.12 strains did.

**Supplementary figure S10.**
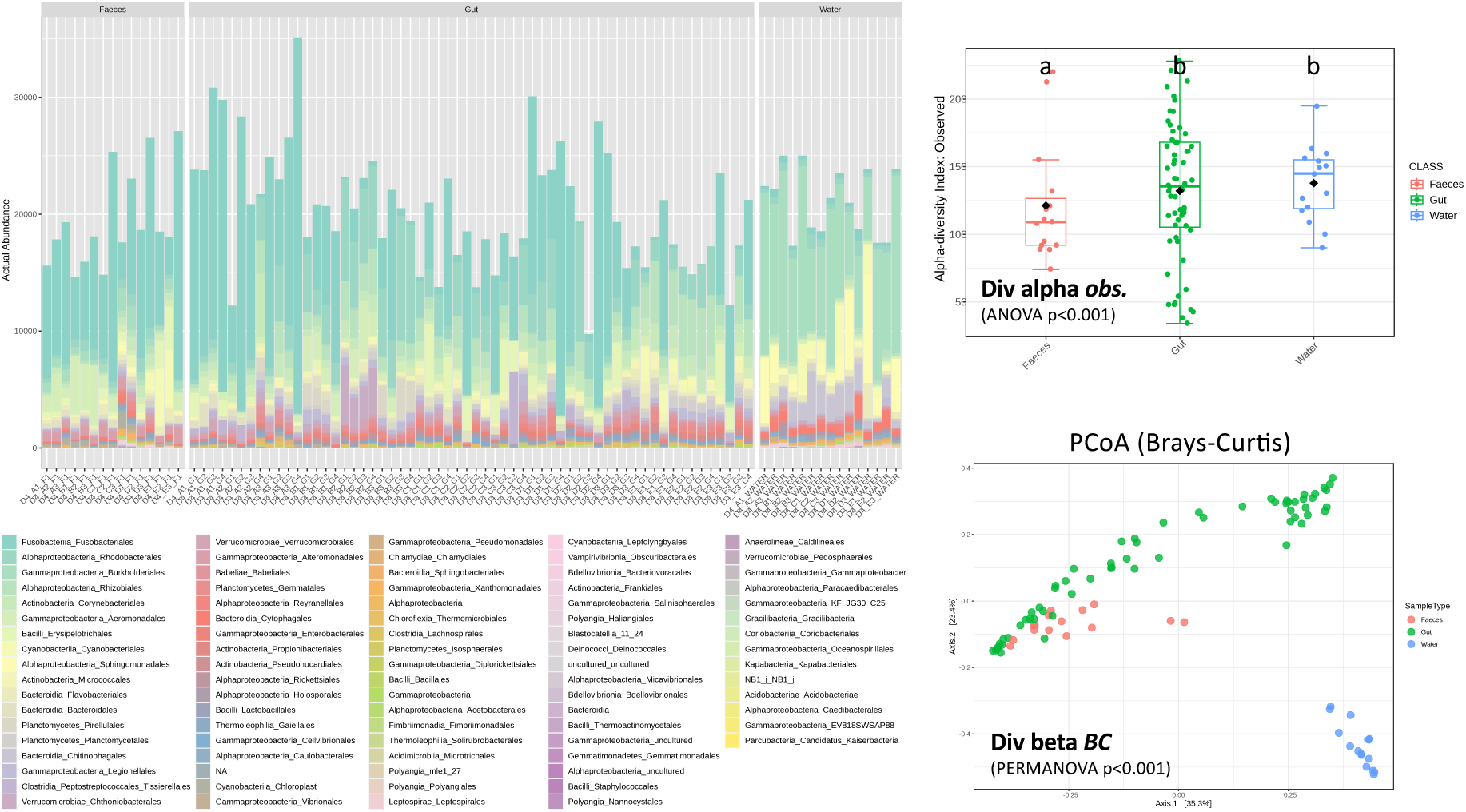
Taxonomic composition of the faeces of the Medaka fish to the different treatment compared to those of the gut microbiote and the water of the different exposure tanks considering the bacterial families and genera levels illustrated individual sample composition histogram (A), shows lower alpha diversity in the faeces (B) and clearly distinct bacterial compositions (Permanova p<0.001) in these 3 biological compartments (C).

**Supplementary figure S11.**
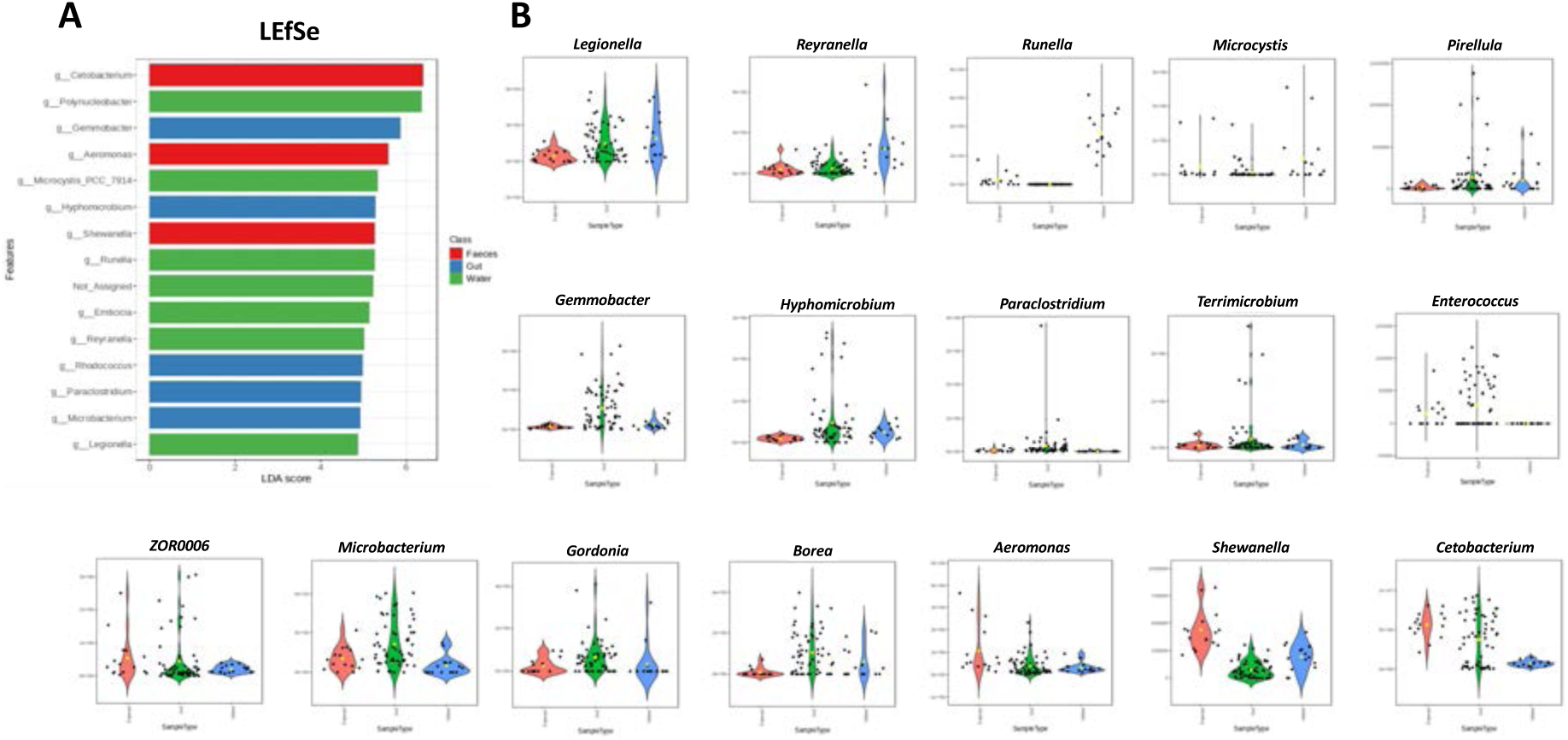
Relative enrichment of the faeces, the guts and the water tanks in certain bacterial taxa illustrated by LEfSe considering all experimental conditions together (A) and boxplots of a selection of bacterial genera presenting heterogenous distribution among these 3 compartments (B), such as *Cetobacterium*, *Shewanella* and *Aeromonas* that appear more relatively abundant in the faeces than in water or in gut bacterial communities.

**Supplementary figure S12.**
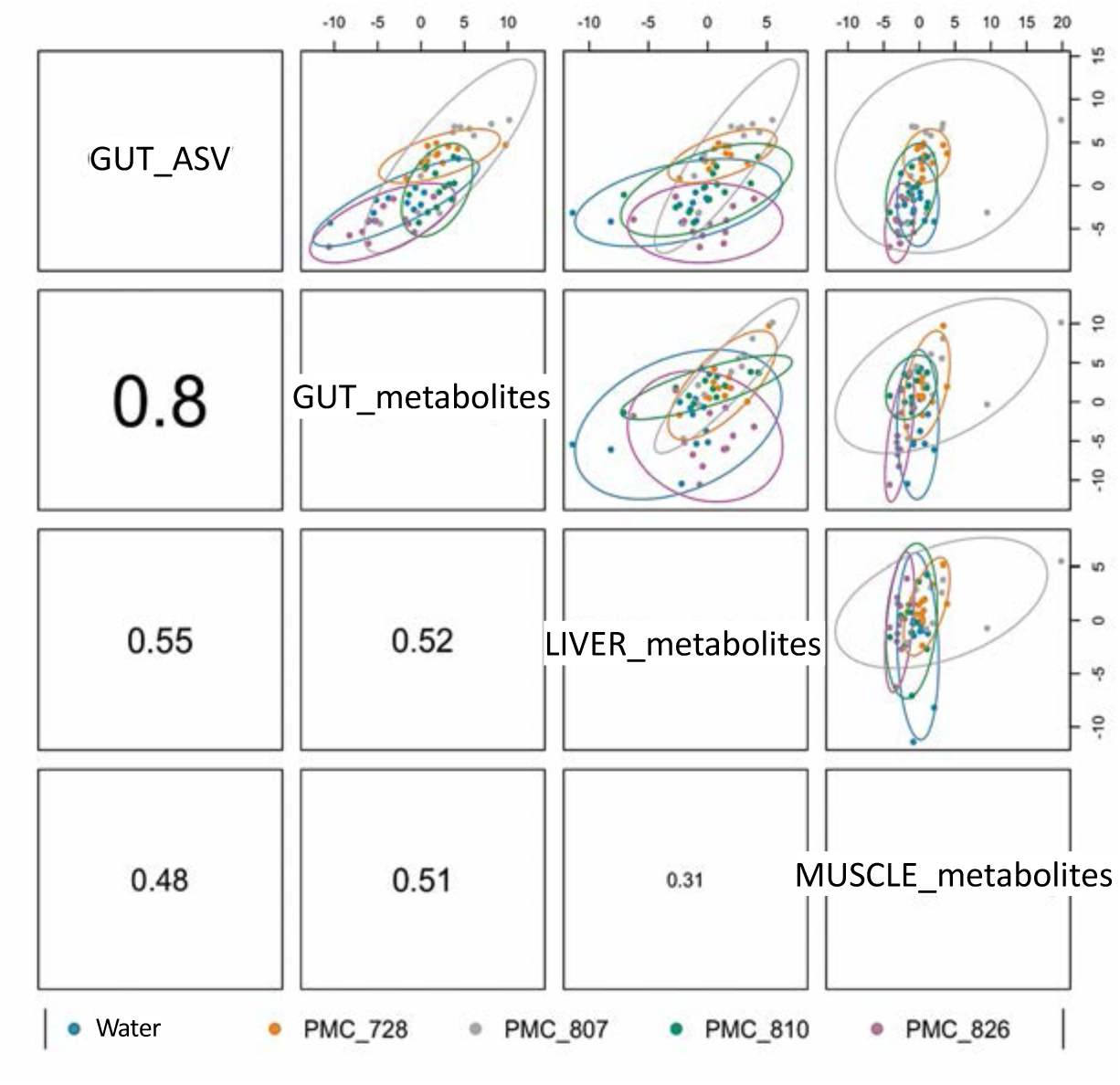
Global correlation matrix of the dataset retrieved from the different analysis performed from the different Medaka fish tissues indicates a better global correspondence (0.8) between the metabarcoding and metabolome analyses performed in the gut.

**Supplementary figure S13.**
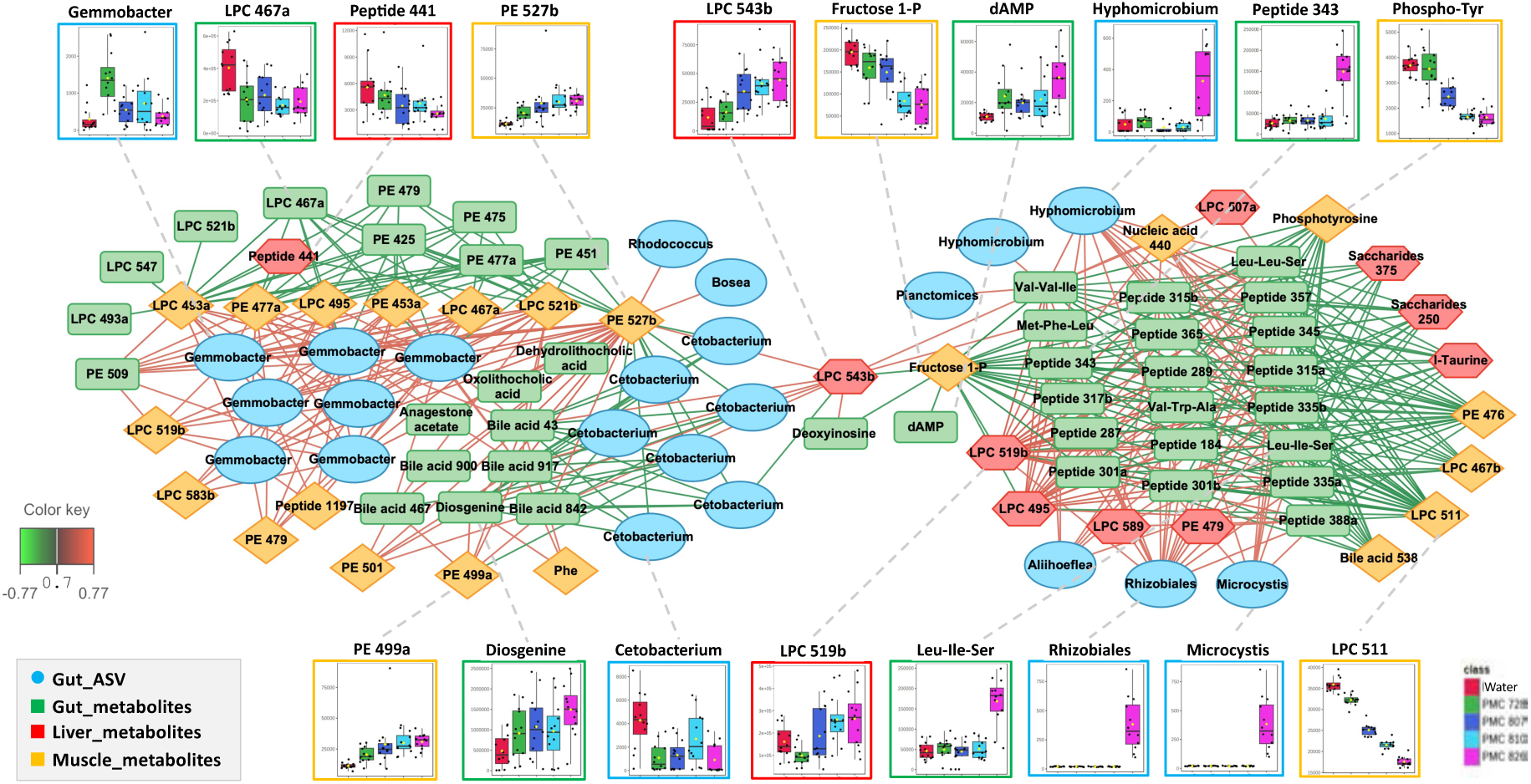
Relevance network analysis performed with MixOmics illustrating the most correlated metabolites (green, red, and orange for gut, liver and muscle metabolome) and gut ASVs (in blue) discriminating among the five treatments considered together. Only variables (ASVs and metabolites) with correlation values above ±0.84, ±0.83, ±0.86, and ±0.92 are displayed, respectively. Pearson correlations between covariate metabolites and ASVs are represented by colored segments (green: negative association, red: positive association).

**Supplementary figure S14.**
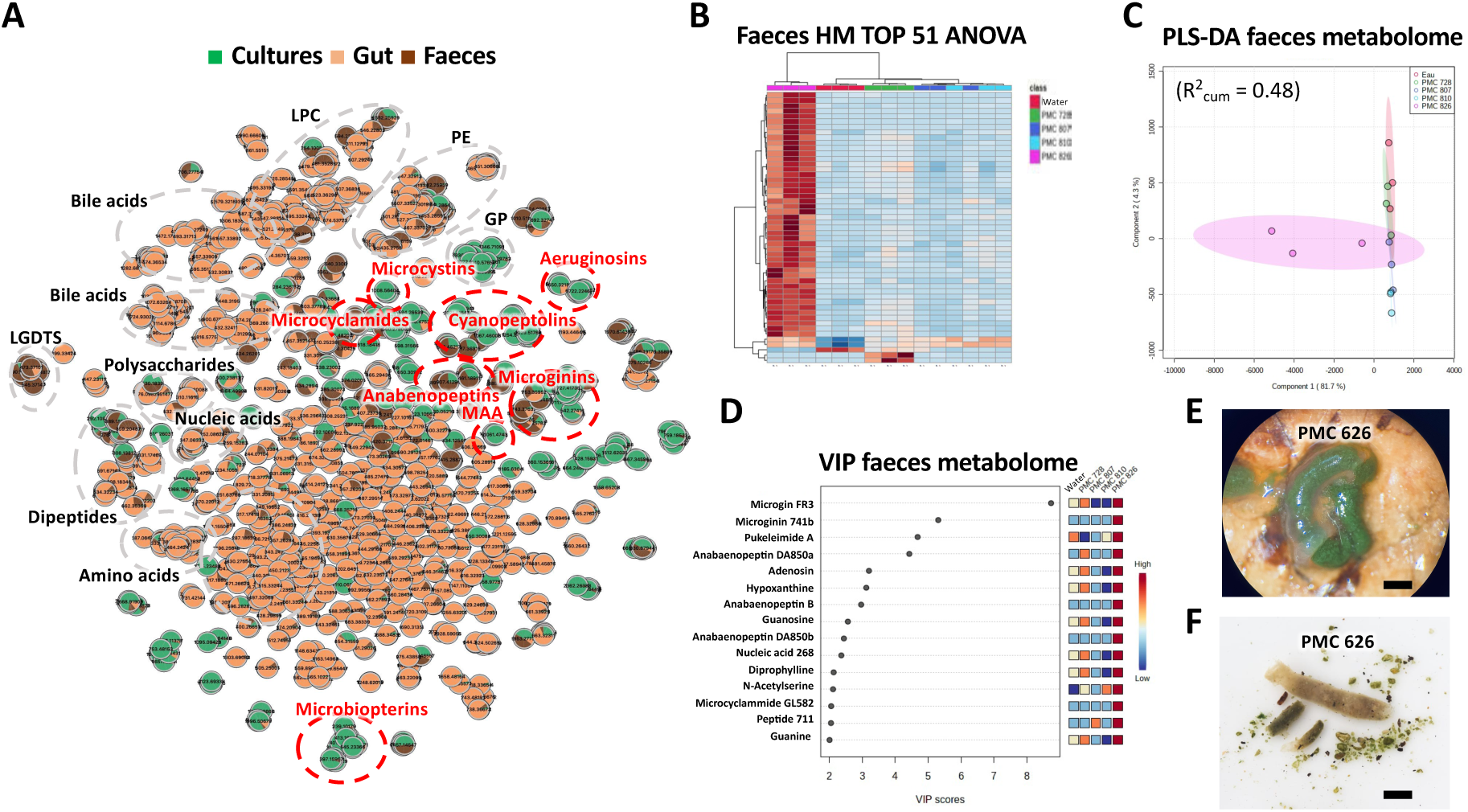
Metabolite composition of the faeces of fish exposed to different treatment collected at the end of the 4-days exposure. Molecular networking of metabolites extracted from the *Microcystis* culture biomass, the gut and the faeces of Medaka fish exposed to the different treatments generated with *t*-SNE algorithm, with cyanobacteria peptide and primary metabolite clusters indicated in red and black, respectively (A). Heatmap with hierarchical clustering of 51 metabolite differentially detected (ANOVA P<0.001) in the faeces of fish exposed to the different treatments (B) and corresponding Individual plot (C) and best VIP (D) of PLS-DA of the faeces metabolomes. Representative pictures of gut (E) and faeces (F) being full of undigested Microcystis of fish exposed to 4-days balneation with PMC 826.12 culture.

**Supplementary figure S15.**
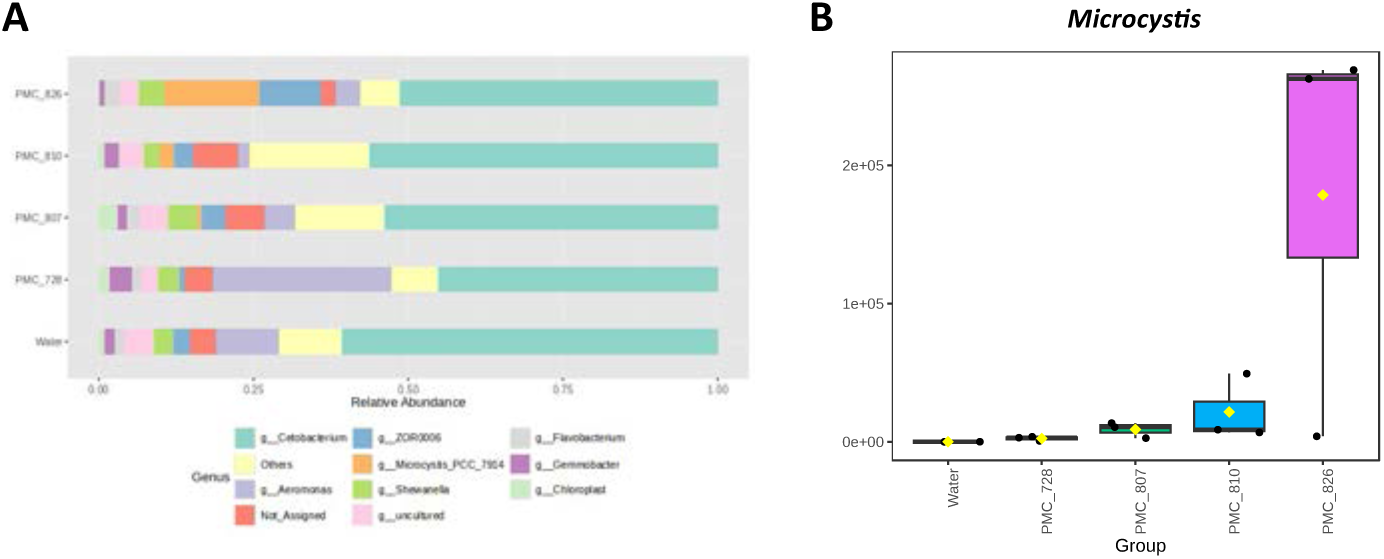
Bacterial composition at the genera level of faeces of fishes exposed to the different *Microcystis* strains (A) and corresponding boxplot of *Microcystis* relative quantification with the different experimental groups (B).

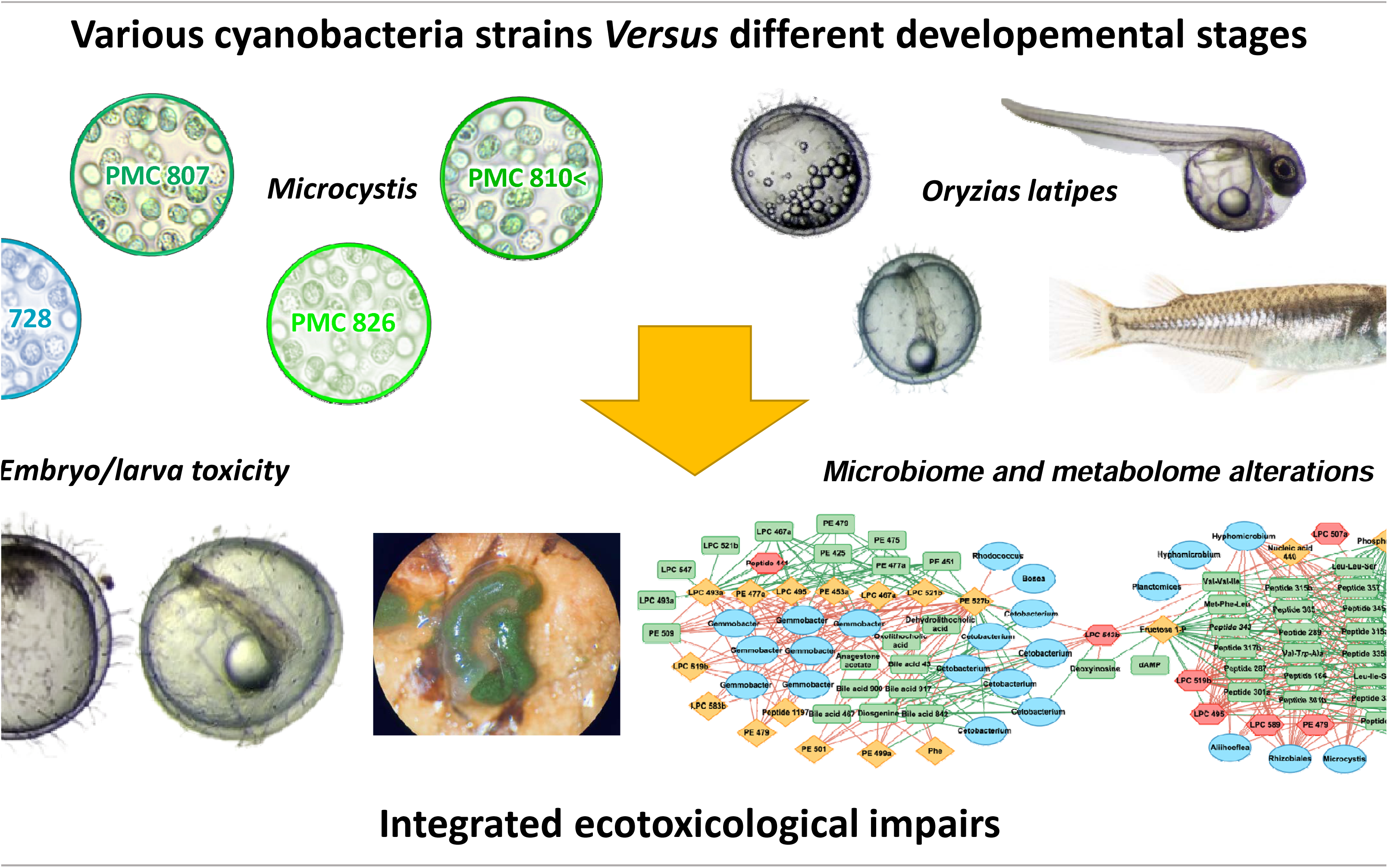

